# Capturing the dynamics of microbiomes using individual-specific networks

**DOI:** 10.1101/2023.01.22.525058

**Authors:** Behnam Yousefi, Federico Melograna, Gianluca Galazzo, Niels van Best, Monique Mommers, John Penders, Benno Schwikowski, Kristel van Steen

## Abstract

**Background:** Longitudinal analysis of multivariate individual-specific microbiome profiles over time or across conditions remains a daunting task. The vast majority of statistical tools and methods available to study the microbiota are based upon cross-sectional data. Over the past few years, several attempts have been made to model the dynamics of bacterial species over time or across conditions. However, the field needs novel views on how to incorporate individual-specific microbial associations in temporal analyses when the focus lies on microbial interactions.

**Results:** Here, we propose a novel data analysis framework, called MNDA, to uncover taxon neighbourhood dynamics that combines representation learning and individual-specific microbiome co-occurrence networks. We show that tracking local neighbourhood dynamics in microbiome interaction or co-occurrence networks can yield complementary information to standard approaches that only use microbial abundances or pairwise microbial interactions. We use cohort data on infants for whom microbiome data was available at 6 and 9 months after birth, as well as information on mode of delivery and diet changes over time. In particular, MNDA-based prediction models outperform traditional prediction models based on individual-specific abundances, and enable the detection of microbes whose neighbourhood dynamics are informative of clinical variables. We further show that similarity analyses of individuals based on microbial neighbourhood dynamics can be used to find subpopulations of individuals with potential relevance to clinical practice. The annotated source code for the MNDA framework can be downloaded from: https://github.com/H2020TranSYS/microbiome_dynamics

**Conclusions:** MNDA extracts information from matched microbiome profiles and opens new avenues to personalized prediction or stratified medicine with temporal microbiome data.

## 1 Background

The human gut is a complex ecosystem where microbes interact amongst themselves and with the host [1]. Dysbiosis, defined as an imbalance in the microbiome of the human gut, has been linked to several complex diseases [2]. It may be reflected by alterations in microbial co-abundance [3] or by changes in how microbes in a community interact with each other. Microbiome interaction (or co-occurrence network) has been shown to potentially exhibit rich information about various health conditions [1]. Disease specificity has been demonstrated for conditions such as inflammatory bowel disease and obesity [3]. Furthermore, the gut microbiome involves a dynamic ecosystem, with microbial co-occurrences or interactions changing over time [4], potentially providing information about health-to-disease transitions [5, 6]. Variations in the human gut microbial ecosystem can be caused by multiple factors, including a sudden change in diet [7] or drug intake, exemplified by metformin [8], a medicine to prevent or treat Type 2 diabetes, and antibiotics [9]. Microbial perturbations may have short- or long-term health effects. For instance, perturbations to the microbiome during infancy have been associated with the development of chronic illnesses in later life, including infectious diseases and asthma or allergies [10, 11, 12].

Two important and highly studied determinants of early-life microbiome establishment are birth mode and infant diet. C-section delivery provides a barrier in the dispersal of maternal fecal and vaginal microbes during delivery and has been linked to various non-communicable diseases and least in part as a consequence of its perturbation in early-life microbial colonization [13, 14, 15]. In addition, the shift from infant feeding (breastfeeding and/or formula) to a more diverse diet consisting of a wide variety of substrates in complementary foods which largely takes place between 6-9 months post-partum, has been associated with a rapid diversification and maturation of the intestinal microbiome [14]

Capturing time-related patterns in data can be achieved via time series analysis (TSA) or longitudinal analysis (LDA). Such analyses involve time course data, and extract additional information from the data compared to cross-sectional studies. The latter involves analysing data limited to a single time point only. The terms time series and longitudinal analyses are sometimes used interchangeably. However, there is a subtle difference, and some methods developed for TSA may not transfer well to LDA contexts. Whereas time series data refer to a sequence of data points, collected at multiple time points or intervals, longitudinal data refer to a subject’s or object’s measurement(s) taken over time. Standard statistical LDA approaches cannot simply be transferred as such to the microbiome field. This is because of the characteristics of metagenomics data. The noisiness, compositionality, and sparseness of the data pose a big challenge in modelling microbial structure; the complexity of multivariate repetitive data for each individual adds to this challenge. In general, methods for microbiome time course data typically aim to address either one or a combination of the following questions (see also [16]). Is there a temporal trend? What is the similarity between multiple time-course profiles? Which microbial community members co-evolve? For examples of microbiome LDA analyses that address such questions, we refer to references in [17].

The construction and interpretation of personalized networks have obtained renewed attention in the context of precision medicine. For instance, Menche et al. [18] used a template network structure derived from knowledge about protein interactions and, for each individual, superimposed the individual’s gene expression scores on nodes. For personalized networks derived in this way, node values are specific to individuals, but edge values are constant across individuals based on reference data. In contrast, we define an individual-specific network (ISN) as a network, for which both nodes and edges can be allocated to a single individual, and that can be seen as a realization of a new measure to describe within-individual activity. An example of an ISN that only uses data from a single individual would be a beta-cell interaction network for beta-cells residing in an individual’s tissue sample and edges defined by changes in intracellular calcium concentrations [19]. One procedure to construct ISNs from reference data was proposed by [20, 21] and has been applied to several scenarios [22]. In [22], the authors compare and evaluate Kuijjer’s LIONESS (Linear Interpolation to Obtain Network Estimates for Single Samples) method [20] and ssPCC (single sample network based on Pearson correlation) [23] for metabolomics measurements and two independent groups of individuals (for instance cases and controls). In ssPCC methodology [23] only the perturbation that an individual causes to an interaction network, when adding the individual to the pool of samples, is considered. In contrast, LIONESS constructs an individual-specific network for each individual of interest from perturbations that subtracting the individual from a pool of individuals causes on the population-based interaction network. Given that the development of individual-specific networks is recent, and given the challenges involved with compositional data, only a few applications exist with microbiome data. For instance, Mac Aog’
sain et al. [24] used LIONESS on microbiome abundances as nodes and Pearson correlations as edge strengths. Individual-specific interactions (edges) are taken as new predictors in models for the time to the next exacerbation in a chronic airway disease. Until now, Individual-specific edges were not used to understand the dynamics of individual-specific microbial interaction profiles. An interesting individual-specific approach to the temporal analysis of microbiome data was introduced by Yu et al [25]. Their method builds on an earlier work, particularly the *individual-specific edge-network analysis* (iENA) framework [26], with disease prediction as the objective. iENA overcomes a critical practical difficulty of the ENA framework: typically, there are not enough longitudinal samples available for the same individual. Their adaption to microbiome data selects microbial interactions (edges) as biomarkers with only a limited number of samples from each individual. Selection is based on a strategy based on edge pairs, and edge strength is based on a form of correlation as a measure of association. An additional application on faecal microbiomes can be found at [27].

For the first time, we here use Kuijjer’s LIONESS method to study microbial dynamic patterns via individual-specific microbiome networks. As explained in the Methods Section, their method relies on reference data, *i*.*e*., a collection of samples that can be pooled to compute a microbiome co-occurrence network. Several strategies exist for constructing a pooled-data microbiome co-occurrence network. These strategies either rely on Pearson-like correlations or on graphical models and the inference of sparse variance-covariance matrices. For an overview and discussion of common co-occurrence network strategies, we refer to [28, 29]. Generally speaking, a microbiome co-occurrence network takes microbial taxa as nodes and evidence for microbial association as edge strength. In the literature, the microbial association is often called “co-occurrence” or sometimes “interaction”. Pooled-sample or population-based models have shown their utility to increase our understanding of underlying characteristics of individuals or to derive personalized predictions [30, 31]. However, from the perspective of personalised medicine, if association networks were available for each individual separately, then descriptions of such networks would readily be individual-specific. Moreover, taking those individual-specific networks as new units of analysis, one would use more information from the data than is typically done. Such analyses may involve association models to understand mechanisms, prediction models to estimate the risk of disease or treatment non-response, identification of endotypes, or, more generally, homogeneous subgroups of individuals that may be targeted together during drug development processes.

In this work, we develop a novel approach to study individual-specific microbial neighbourhood dynamics over time or across conditions. We illustrate the approach on microbial data from newborns with measurements at 6 and 9 months over time. We first describe the study design, and the components needed to compute individual-specific microbiome networks. Second, we introduce a new microbiome analysis framework, based on representation learning, that we call *multiplex network differential analysis* (MNDA). MNDA generates new representations of local microbiome interaction neighbourhoods that can be used in supervised or unsupervised models, or to identify stable or unstable microbial taxa over time or across interventions. Finally, we present and discuss the results of various microbiome dynamic analyses via MNDA. These analyses broadly fall into three broad classes. The first class covers dynamic analyses of global networks, where a global network refers to a microbiome co-occurrence network constructed on a pooled set of individuals. These individuals may share the point in time at which data are considered for microbiome analysis, or some other—for instance, clinical—characteristic. The second class captures dynamic changes of microbiome ISNs. These analyses include comparing fluctuations in microbiome ISNs over time and tracking each individual in an embedding space. We use microbial neighbourhood dynamics to identify subgroups of individuals that are similar in terms of their microbial interaction dynamics, which provides a different viewpoint than the classical *Dirichlet multinomial mixture* (DMM) clustering [32]. The third class of analyses aim to enhance the interpretation of findings: topologies of microbial association networks are studied within and between individuals and are linked to infant delivery mode and infant’s diet changes between months 6 and 9.

## 2 Methods

### 2.1 Study Design and Microbiome Profiling

The LucKi Gut cohort is an ongoing study monitoring gut microbiota development throughout infancy and early childhood [33]. Pregnant women from the South Limburg area in the Netherlands were recruited via mother and childcare professionals, through the study website and through Facebook. Women were eligible to participate if they gave birth at *>* 37 weeks of completed gestation. Study questionnaires and faecal samples of the infant were collected at different time points, e.g., 1-2 weeks, 4 weeks, 8 weeks, 4, 5, 6, 9, 11 and 14 months. Parents were instructed to collect infant faeces from diapers and freeze them immediately at –20^*◦*^*C* in their home freezer inside a cool transport container (Sarstedt, Hilden, Germany). Samples were transported to the laboratory, preserving the cold chain. Metagenomic DNA was extracted with a custom extraction protocol involving mechanical and enzymatic lysis [34]

Microbiome profiling was performed by next-generation sequencing of the 16S rRNA V3-V4 hypervariable gene region. Thereafter, a DADA2-based pipeline was used to identify Amplicon Sequence Variants ASVs [34]. Lastly, a *centered log ratio* (CLR) transformation of data using the ALDEx2 R package was performed to account for the compositional nature of microbiome data, whenever appropriate [35]. Additional details are provided in earlier publications [36].

The current study focuses on months 6 and 9 after delivery. These moments in an infant’s life are recognized milestones in the maturation of microbial communities, possibly influenced by change of diet during 6-9 months after birth. For this reason, we include dietary information on infants in applications of our new framework. Diet type was encoded as *{*0, 1, 2*}*, with “0” representing breast milk exclusively, “2” representing exclusively solid food, and “1” indicating a mix of both. We refer to a diet as *persistent* if it does not change during 6-9 months. Furthermore, we also had information about the mode of delivery (either C-section or vaginal delivery) available at months 6 and 9. We used this information in prediction models and stratified analyses.

### 2.2 Data Pre-processing and Exploratory Analysis

Selecting informative individuals and taxa and filtering out random noise was achieved with the following prevalence filter: only amplicon sequence variants with a prevalence exceeding 15% survived the filtering. Prevalence indicates the percentage of samples in which a microbe was detected. The average sequencing depth was 57392 read counts, while the range was [11123, 105921], with an interquartile range of 25346. Pre-processing was done on the merged data of 155 newborns across months 6 and 9. No infants were removed. Out of 1144 taxa, only 95 (8%) remained after data pre-processing. These 95 taxa were considered for subsequent analyses. In addition, we defined two new classes of microbes: *appearing* or *disappearing* microbes – taxa that meet the 15% threshold at month 9 (appearing), or at month 6 (disappearing), but were filtered out by joint-time point pre-processing.

Basic exploratory analysis involved computation of *α-diversity* (within-sample diversity), at each time point. In the spirit of a CoDA analysis, we used *Aitchison’s distance* (Euclidean distance on CLR-transformed abundances), as implemented in the *vegan* R package [37]. We also investigated *β-diversity* or between-sample diversity. In particular, CoDA ordination was implemented via PCA on CLR-transformed compositions. We used the implementation of Le Cao et al. [38]

### 2.3. Global Microbial Network Construction

For the remainder of this manuscript, we refer to a *global network* as a network computed over a set of independent individuals. For instance, such a set can refer to all infants at month 6 after birth. We computed a global network for each time point, by selecting microbial taxa as nodes and calculating association strength via *microbial sssociation graphical model analysis* (MAGMA) [39] on all newborns at each timepoint (81 at timepoint 6m and 74 at 9m). MAGMA uses *copula Gaussian graphical models* for which marginals are modeled with zero-inflated negative binomial generalized linear models, and sparseness is induced via a graphical Lasso strategy. From a practical point of view, we used the *rmagma* R package to derive MAGMA-networks (https://gitlab.com/arcgl/rmagma). To optimize its internal penalization parameter, we used the rotation information criterium (RIC) [40]. For more details about the adopted MAGMA analysis, see Additional file 1 – *Global Network Construction*.

We preferred MAGMA over commonly used correlation-based measures due to its theoretical advantages [38] and its flexibility to adjust for confounders. In the case of microbiome data, characterized by zero-inflation, high heterogeneity and overdispersion, Pearson correlation as a measure of association may give rise to false positives, as shown by Friedman and Alm 2012 [41]. For an overview of microbial co-occurrence network methods, we refer to [29]. Notably, MAGMA was not included in this mini review. MAGMA has the advantage of yielding a sparse microbiome co-occurrence network, acting as a natural sparsifier. It targets direct associations, removing indirect ones. As illustrated by Kishore et al. [42], there are remarkable differences in microbial co-occurrence networks across inference methods. We also selected *SparCC* [ref https://doi.org/10.1371/journal.pcbi.1002687 same as before!]as an alternative to MAGMA, to assess the impact on conclusions of the selected microbial co-occurrence network inference method. Even though SparCC can better handle spurious associations compared to its predecessor correlation-based microbial cooccurrence network inference methods, it may fail to generate a positive definite covariance matrix. In practice, we applied *FastSpar’s C++* implementation. The microbiome dynamics analysis results with SparCC, parallel to MAGMA, are presented and discussed in Additional file 3 – *SparCC Analyses*. We emphasize that pre-processed data entered MAGMA and SparCC analyses with default options. No data transformation was performed prior to MAGMA/SparCC, as those analyses frameworks internally accommodate compositional data.

### 2.4 Individual-Specific Network Construction

We used Kuijjer et al.’s LIONESS method [20] to infer individual-specific networks from global microbiome co-occurrence networks, as implemented in [21]. Each individual-specific edge weight measures the impact of the individual observation on a global network edge. In particular, the edge weights of the *n*_*th*_ ISN (for individual *n*) were computed as

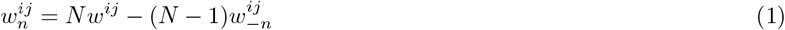

where *w*^*ij*^ and 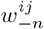 are the edge weights of a global network (Section 2.3) and the *n*_*th*_ leave-one-out network, respectively, for any pair of microbes (*i* ≠ *j*). *N* is the total number of individuals in a reference population (Figure 1). For large populations (*N* → ∞) and under a weight homogeneity condition, the average individual-specific edge weights 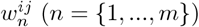 (*n* = {1, …, *m*}) converges to the corresponding global edge weights. Namely, the global network can be seen as a weighted average of the ISNs. Weight homogeneity means that the proportion of the weights between individuals is constant between global (*w*^*ij*^) and LOO 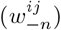 networks ([20] Suppl. 5.2). We considered our 69 paired infants at month 6 and at month 9 after birth as two distinct populations. We do that by only considering the paired infants in both timepoints. Since in this work we are only interested in unidirectional strengths of microbial associations, we replaced all individual-specific edge weights with their absolute values. Notably, even when population-based edge weights, *w*^*ij*^ and 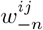, are positive, the derived individual-specific weight 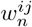 may be negative. This occurs when 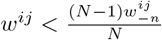

**Figure 1.**
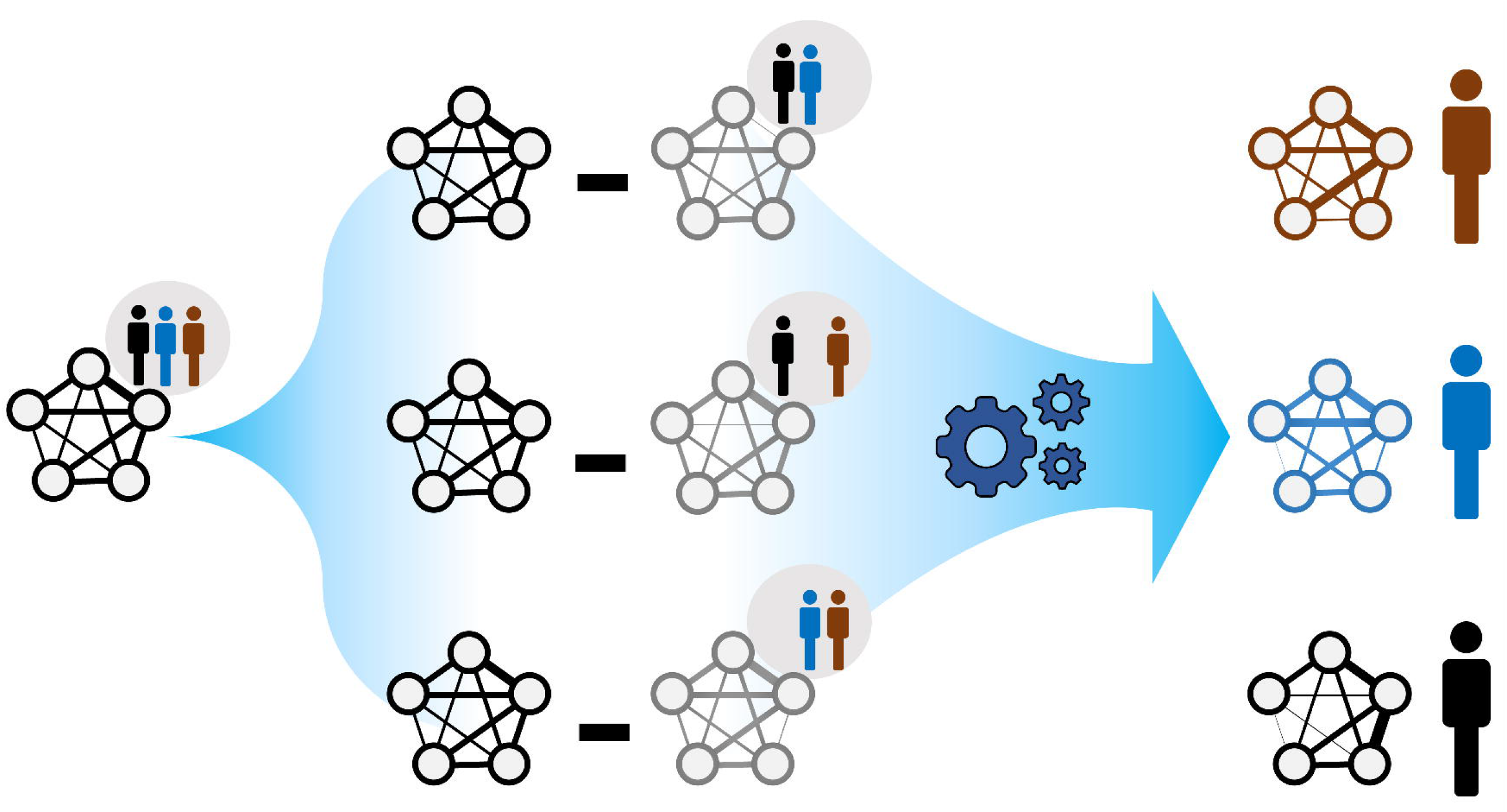
The rationale behind ISNs. A global network summarises node and edge information across the individuals belonging to the same population. The goal is to compute individual-specific networks with individual-specific edge weights (as in [21]; each individual is allowed to exhibit a particular network topology. Comparing such network topologies may not only reveal individual heterogeneity, but may also indicate instances where health-related interventions based on the global network may be inappropriate.

### 2.5 Multiplex Network Differential Analysis

The ISNs constructed in the previous section can be paired into 69 multiplex networks (*i*.*e*., networks with multiple layers of matching nodes [43, 44]). Each multiplex refers to a single individual; within each multiplex, a layer refers to an individual-specific network at a particular time. More generally, we assume as input to a representation learning algorithm, a number of multiplexes that represent an object, for which matched data are available. Matching may be performed on the basis of repeated measures over time for the same individual (as it is the case for our study example), but it may also refer to matched samples where two individuals are matched according to shared characteristics. In both scenarios, the availability of individual-specific edges offers the opportunity to investigate local neighbourhood stability.

To capture how the local neighbourhoods of microbial taxa change over time, we proposed the following algorithm, which we term *multiplex network differential analysis* (MNDA). First, multiplexes were formed. For our LucKi subcohort, ISNs derived in Section 2.4 at months 6 and 9 are paired. Each ISN has 95 nodes, representing the 95 microbial taxa retained in the study, after data pre-processing (Section 2.2). One individual and its multiplex ISN structure is shown in Figure 2A. Second, we developed a network representation algorithm, based on a shallow *encoding-decoding neural network* (EDNN) that forms the core of the MNDA framework (Figure 2B). For the implementation, we used the *Keras* R package. A graphical flow is given in Figure 2B. The inputs and outputs to the EDNN are vectors, one for each node in a layer of a multiplex. The input vector at the encoder side uses the edge information of each node’s immediate (graph distance 1) neighbours. A binary such vector for a particular microbial taxon for individual 1 at months 9 would be a vector of ones and zeros indicating which are the direct neighbouring taxa based on the individual’s ISN at month 9. This assumes a binary ISN (*i*.*e*. an unweighted network with edges that are either present or not). It is worth mentioning that, in this method, we do not consider self-loops (*i*.*e*. the node itself is not seen as a direct neighbour of itself), which makes our model generalizable to accepting new nodes. A non-binary (weighted) input vector would be a vector for which the ones in the binary vector are replaced by their actual edge strengths. It is here that an additional argument can be made in favour of microbiome cooccurrence network inference methods that lead to sparse networks. MAGMA will generate a sparse weighted network, avoiding the need to work with fully connected networks. Fully connected networks may complicate the interpretability and are less computationally tractable than sparse networks.

**Figure 2.**
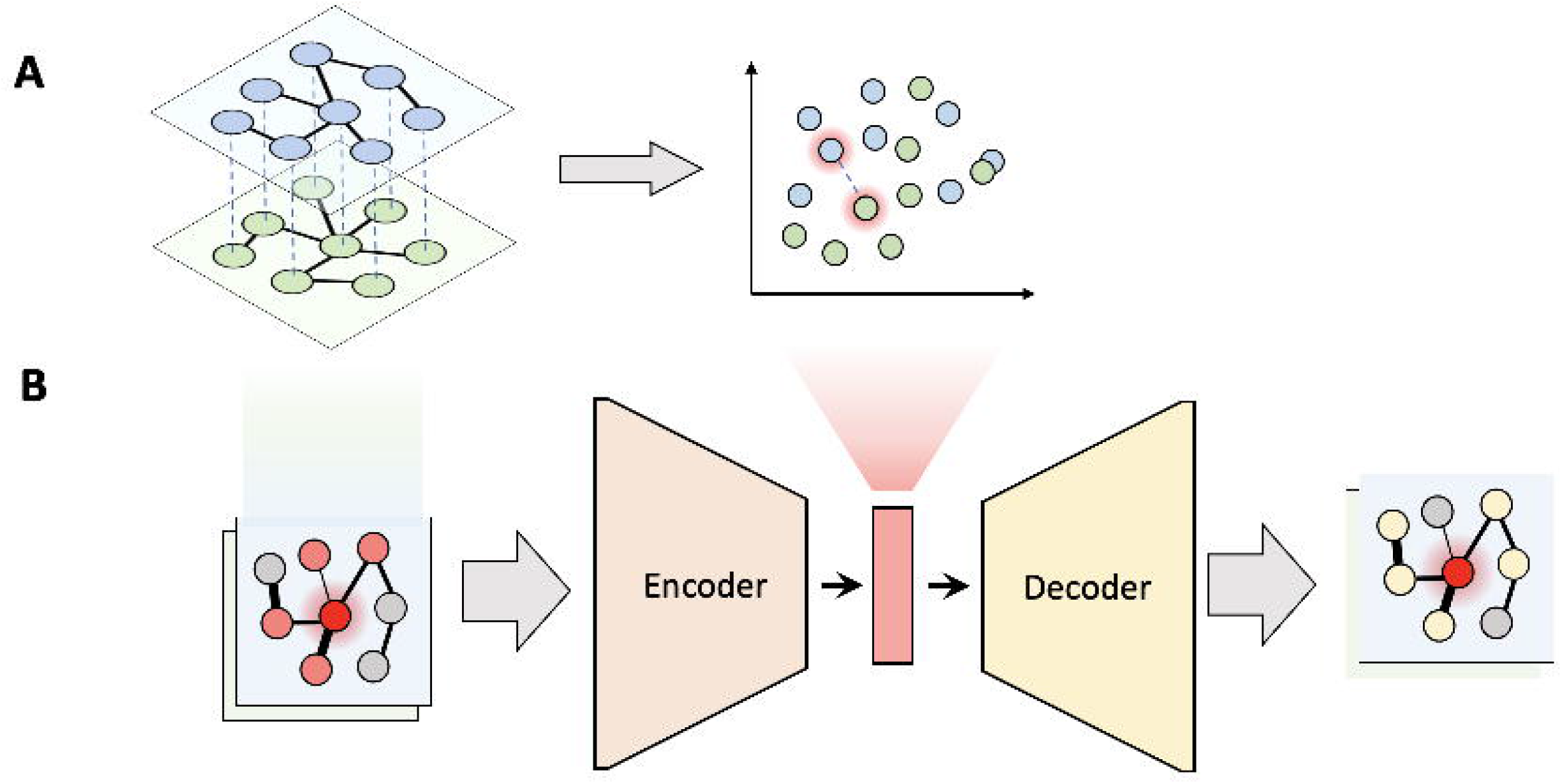
Multiplex network differential analysis (MNDA) framework. (A) in multiplex network representation learning, all the nodes of a multiplex network are transformed into an embedding space; the highlighted nodes are matched pairs and, in our case, correspond to the same microbe. (B) Multiplex network representation learning is performed using an encoder-decoder neural network for all individuals.

The output vector of EDNN, what needs to be predicted at the decoder side, is a representation of more distant neighbours likely to be reached by a *random walk*. In particular, for a specific (seed) taxon, both binary or weighted versions of such an output vector would refer to probabilities that a microbial taxon is reached by a *fixed-length random walk* starting from the seed taxon. Binary and weighted versions differ in the way these probabilities are computed (see next – *A customized implementation of repetitive weighted random walks*). As activation functions for the hidden neurons and output neurons, we used *ReLU* and the *Logistic* functions, respectively. The dimension of the hidden space is equal to the number of hidden units, and was chosen to be equal to 10 since this resulted in the least *mean squared error* of EDNN compared to the other choices (i.e. 2, 5, 10, 15, 20). Third, after having learned the local structure of multiplex network layers and having created representations in a 10-dimensional embedding space, we tracked the positions of the same microbial taxon at months 6 and 9. We formed these pairs for all 95 taxa (Figure 2A), and do this for every individual. We then computed a distance between paired taxa (see next – *A new measure of microbial dynamics*).

#### A customized implementation of repetitive weighted random walks

To obtain the node visit probabilities at the decoder side of our MNDA framework, we developed a *fixed-length weighted random walk algorithm*. Accordingly, the walker is restricted to take a limited number of steps to a predefined fixed length. This enables us to characterise the local structures of the ISN. More specifically, we start “walks” from each node and compute the probability of visiting all the other nodes during the walk. We tried different values for walker length (*≤* 10) to keep the walks local. We finally chose a walk length of 5 that resulted in the smallest mean square error for EDNN. It is worth mentioning that the results were fairly stable across walk lengths. To calculate the node visit probabilities, we repeated the walking process 100 times for each node.

Since the degree of co-occurrences of microbes may be highly variable, we proposed to use a *weighted random walk algorithm* to take the edge weights into account. To move from node *i* and determine the next step in a simple random walk algorithm, all the neighbouring nodes have the same probability of visit 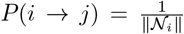, where *N*_*i*_ is the set of neighbours (distance 1) of node *i* with operator ∥.∥ counting its members. In the weighted version of this algorithm, the probability of moving from node *i* to node *j* is proportional to the edge weight *w*_*ij*_ linking node *i* to node *j*:

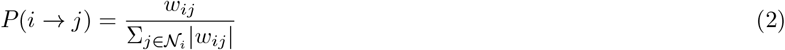

Although the idea of weighted random walks is not new, to our knowledge, no customized code was available for their use in our framework. Our code is provided as part of a *GitHub* repository that covers the workflows used in this manuscript (https://github.com/H2020TranSYS/microbiome_dynamics).

#### A new measure of microbial dynamics

The low-dimensional learnt local structure of the multiplex network layers can be analysed further by computing the angle *θ* between embedded vectors of belonging to the same microbial taxon at different time points, or the corresponding *cosine similarity cos*(*θ*). In a positive space, the smaller 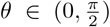, the larger *cos*(*θ*) and thus the larger the similarity between vectors. Even though it is not a genuine distance metric, the cosine distance *d*_*cos*_(., .) is often used as a complement of cosine similarity in a positive space, and defined as 1 − *cos*(*θ*) (without any restrictions on *θ*):

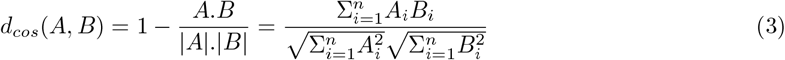

with *A*_*i*_ and *B*_*i*_ (*i* ∈ {1, …, *n*}) components of *A* and *B*, respectively, and *A, B* time-specific representations of the same microbial taxon as vectors in the derived embedding space. In contrast, the normalized angle between *A* and *B* (called angular distance) is a metric but is defined as 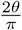 in positive space and 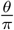 else. As cosine similarity varies in the range [−1, 1], cosine distances vary in the range [0, 2].

#### Simulation study

We conducted a simulation study to evaluate the ability of MNDA to capture the neighbourhood variation in a multiplex network. To this end, we constructed a graph with node degree and weight distributions similar to our microbiome graphs. Then we created a perturbed copy of this graph with a few (*m*) distinct randomly selected (“control”) nodes having different neighbouring nodes. This resulted in a multiplex network with control nodes whose neighbourhood varies. We expect that in the embedding space all the nodes are very close to each other except for the control nodes. Additional file 3: Figure S7a shows the embedding space for a simulated 2-layer multiplex network; observations are in line with the aforementioned expectation. To quantify this observation, we first sorted the cosine distance between the node pairs and select the m most distant pairs. Next, we calculated the *Jaccard index* between the pre-set varying nodes and MNDA-based detected nodes and observed that it almost always equals one. Hence, our proposed measure can, with a high degree of confidence, detect nodes whose neighbourhoods change from one graph to the other. To further assess the robustness of our method under the noise, we added uniformly distributed noise, ranging between *min* = 0 and *max* ∈ {e^−10^, …, *e*^−2^}, to the adjacency matrix of each graph. The Jaccard index was calculated and plotted against different noise levels in Additional file 3: Figure S7b (red diagram).

As for comparison, we used a method based on the eigenvectors of the *Laplacian matrix* of the graphs, which is a typical approach to measure the distance between graphs. We represented the nodes of each graph layer to the eigenvector space of their Laplacian matrix and used the cosine distance between the node pairs. Compared to MNDA, only a few varying nodes were identified by this method (Jaccard index = 0.3). To investigate the robustness of MNDA against random perturbation or noise in the edge weights, we added uniform noise with different ranges to the adjacency matrices of each layer and assessed its performance in prioritising the dynamics. As shown in Additional file 3: Figure S7, MNDA is reasonably robust to such uniform noise and consistently performs better than the eigen decomposition-based method.

## 3 Results

### 3.1 Exploratory Data Analysis

In general, it is well known that the dominant gut microbial phyla are *Firmicutes, Bacteroidetes, Actinobacteria, Proteobacteria, Fusobacteria*, and *Verrucomicrobia*, with the two phyla *Firmicutes* and *Bacteroidetes* representing 90% of the adult gut microbiota [45]. Moreover, the most prevalent phylum for our retained 95 microbial taxa across 6 and 9 months is *Firmicutes* (53 out of 95). *Bacteroidetes* (14), *Actinobacteria* (15), and *Proteobacteria* (13) are (almost) equally represented. *TM7* and *Verrucomicrobia* phyla are represented by a single microbe each (Figure 3). The fractional relative abundances differ between time points. *Actinobacteria* is the most abundant phylum at m6, while *Firmicutes* has the greatest share at m9. Fractional relative abundances per phylum, order, class, and family are provided in Additional file 3 – Figure S1 A–D, respectively.

**Figure 3.**
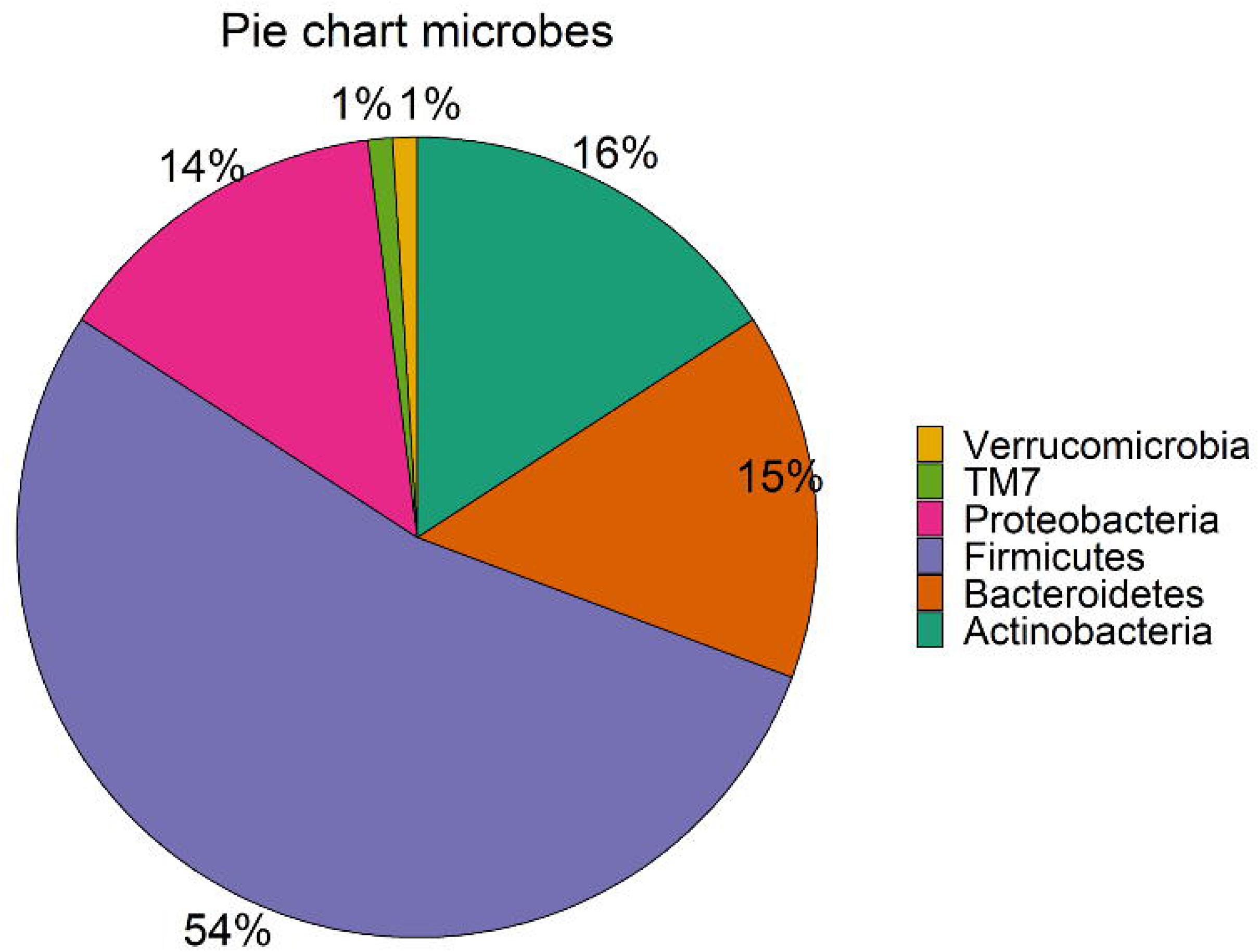
Phyla in the subset of the LucKi cohort. The share of the phyla in the 95 selected microbes is showed. Sample figure title

Microbiome diversity was assessed via *α*- and *β*-diversity on the 69 paired newborns. Violin plots of *α*-diversity grouped per timepoint, delivery type, and diet are depicted in Figure 4. Violin plots combine classical box plotting with kernel density graphs. We found evidence for a significant difference in *α*-diversity between months 6 and 9. Specifically, the paired Mann-Whitney U test shows a low *p-value* (3.843 ∗ 10^−8^), rejecting the hypothesis of no difference between the timepoints. This is in agreement with figures 6A and B, where we can see the increased connection strength at timepoints 9m compared to 6m. Figure 5 shows CoDa ordination plots grouped per timepoint, delivery type, and diet. These plots do not exhibit marked differences between different mode of delivery or diet, while the first principal component provides separation between the timepoints.

**Figure 4.**
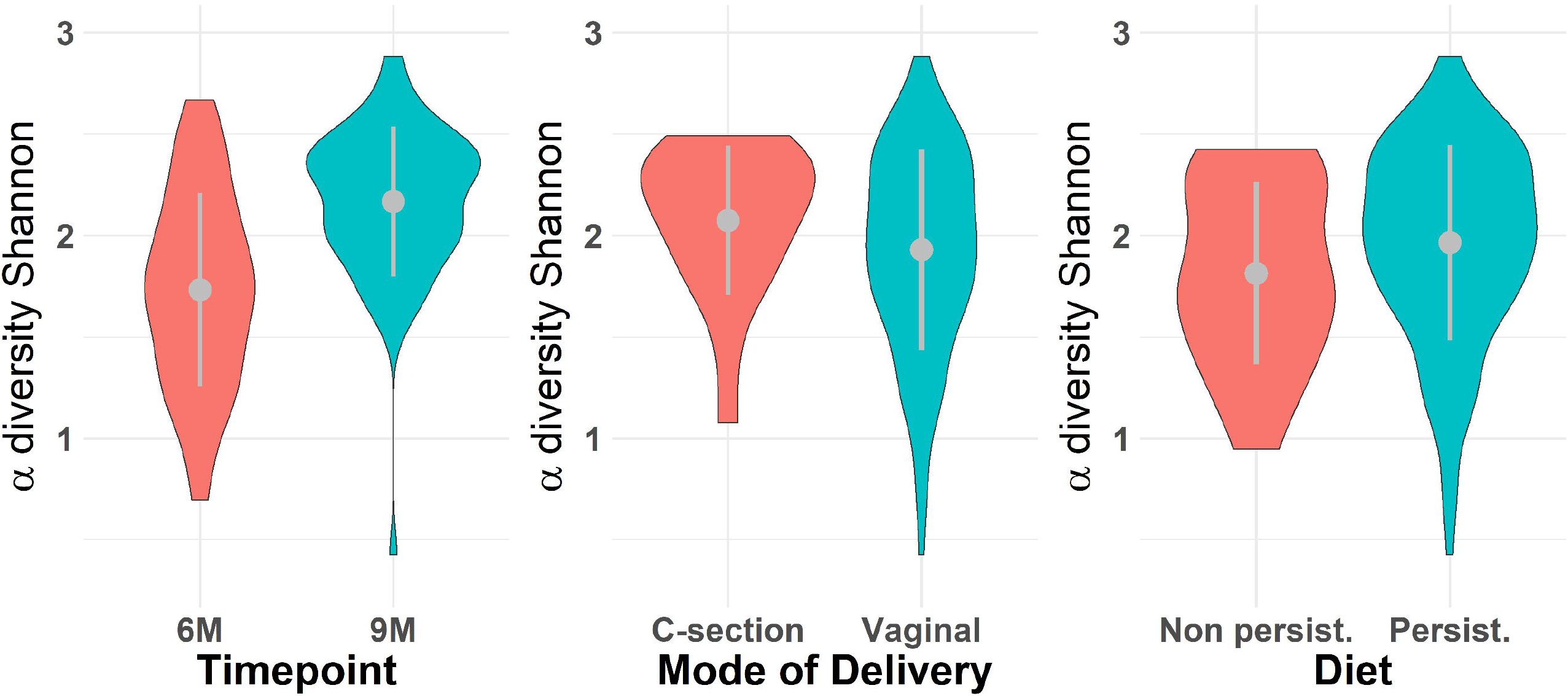
Violin plot for *α*-diversity distribution grouped (A) per timepoint, (B) mode of delivery and (C) diet. The grey dots represent the averages. Gray line extremes indicate plus and minus one standard deviation. Paired Mann-Whitney U test between 6m and 9m rejects the hypothesis of no difference between the timepoints (*p* − value = 3.418 × 10^−8^), while there is no significance (p-value ¿0.05) grouping per mode of delivery and diet. The IQRs are respectively (A) 0.72 for 6m and 0.42 for 9m; (B) 0.45 per C-section and 0.67 per Vaginal; (C) 0.80 per Non persistent and 0.63 per Persistent diet.

**Figure 5.**
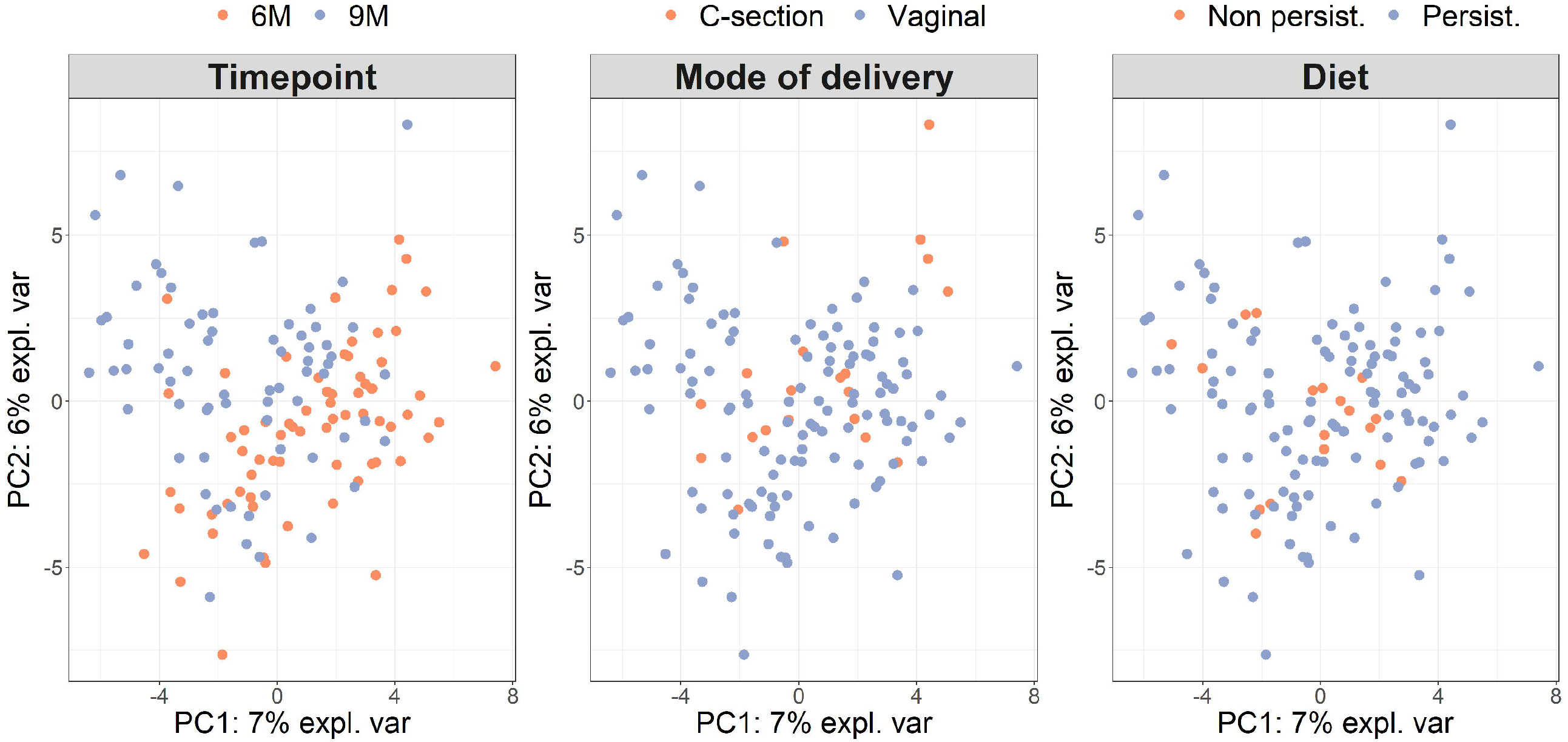
*β*-diversity grouped (A) per timepoint, (B) mode of delivery and (C) diet on paired samples. As in Le cao et al [72], we compute the *β*-diversity with a PCA from the mixOmics package on the CLR transformation of the microbiome data. The axes are the first two principal components and account for 13% of the total variance. Individuals with missing phenotypes (1 for diet) are excluded from the analysis

**Figure 6.**
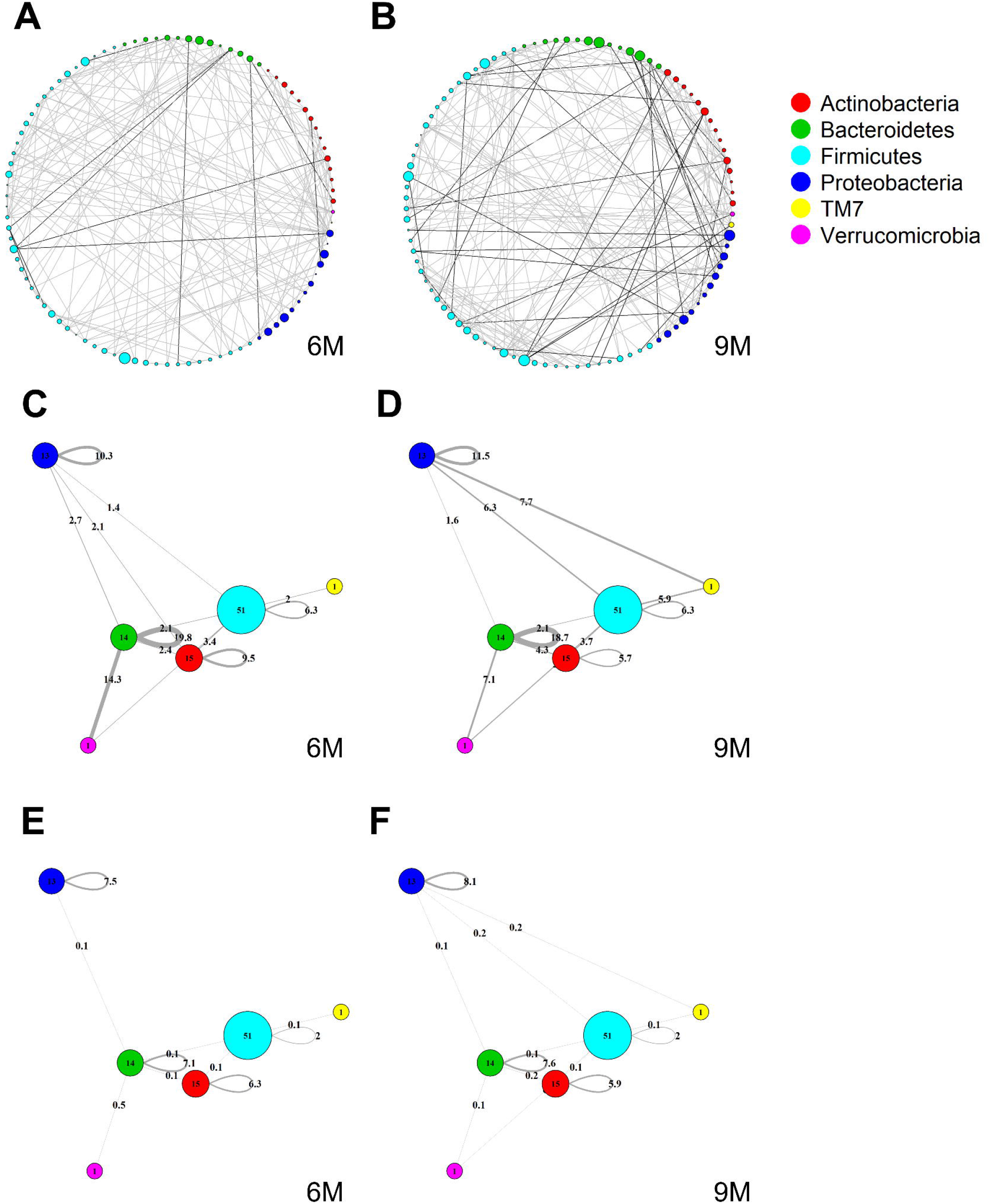
Global MAGMA networks calculated on all subjects retained in the LucKi cohort at 6 (81 subjects) and 9 months (74 subjects) after birth, i.e., available cases at each timepoint. Color code corresponds to phylum classification. The thickness of an edge corresponds to the strength of association. Microbial taxa are organized on a circle according to phylum membership at timepoint 6m (A) and 9m (B). In (C) and (D), respectively timepoint 6m and 9m, edges are aggregated per phylum and within-and across- phyla co-occurrences are computed via sparsified MAGMA binary networks at each time point. Node size and edge weight correspond to respectively phylum size and association strength. In (E) and (F) weighted MAGMA edges are used, whereas binary edges are used in (C) and (D).

### 3.2 Cross-Sectional Global Network Analyses

Figure 4A and B show two global microbiome networks obtained via MAGMA, one per time point, with edge strengths replaced by their absolute values. Microbial taxa are organized according to their phyla. Additional file 2: Figures S1–S3 show similar plots but with microbes organized (coloured) according to order, class, and family, respectively. The size of the node is proportional to relative microbial abundance. The stronger the edge strength, the thicker the edge line that connects corresponding nodes in these networks. Notably, these networks are not fully connected. This sparsification is due to the graphical Lasso in the MAGMA microbial network computation.

Stronger absolute correlations can be observed in the global MAGMA network at month 9 compared to month 6 (Figure 6A). In Figures 6C–F, we grouped microbes of the same phylum into a meta-node and depicted the connections based on the global network co-occurrences. Hence, a node represents a phylum, with a size related to the number of taxa in the phylum, with the exact number indicated in the node itself. The microbes for each phylum can have connections with microbes of the same phylum (indicated by self-loops) or with microbes in different phyla. Edge thickness for edges between phyla is related to the strength of phylum-phylum associations, normalized to account for potential differences in phylum sizes. In particular, for unweighted MAGMA edges (Figure 6C,D), the number of edges *η* between two phyla *P*_1_ and *P*_2_, with respective sizes *n*_1_ and *n*_2_, was benchmarked against the maximum number of edges *n*_1_*n*_2_:

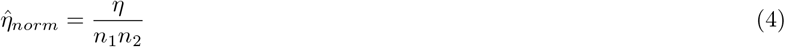

For phylum self-loops, *e*.*g*., in a phylum *P* of size *n*, the denominator in (4) was adapted to 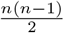. This normalized count has a natural interpretation of a percentage. We obtained the unweighted MAGMA network by binarizing the MAGMA output, with a 1 for every non-zero entry. For weighted MAGMA edges (Figure 6E,F), the connection strength *η* between two phyla *P*_1_ and *P*_2_ was defined as a normalized sum of edge weights, with the same normalizing factor as for binary networks. Note that in this scenario, the interpretation of 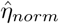as a percentage is no longer possible, since *η* is not normalised for its maximum value.

*Firmicutes* is the largest phylum, with 51 microbes, and shows limited intraphylum and inter-phylum associations. *Proteobacteria* and *Bacteroides* are medium-sized phyla, with 13 and 14 taxa, respectively (Section 3.1), and have strong intra-phylum connectivity. In particular, intra-phylum association strength for *Bacteroides* is close to 20% in the binary global networks (Figure 6C,D) in both timepoints, meaning that almost one-fifth of all the possible *Bacteroides* interactions are present. The strongest binary associations are inter-phylum, appearing at month 9 (between *Firmicutes, TM7*, and *Proteobacteria*). Only one microbe (lowest annotation: Class *TM7-3*) belongs to the *TM7* phylum, with a moderate – and decreasing in time – association with the *Actinobacteria* phylum. Similarly, one microbe (*Akkermansia muciniphila*) belongs to the *Verrucomicrobia* phylum and shows increased connections with *Firmicutes* and *Bacteroides* phyla at 9m. While these phyla associations may be strong in binary global networks, their corresponding association strengths in weighted global networks remains rather limited.

The corresponding global microbiome networks obtained with SparCC are included in Additional file 3: Figure S4 and S5. Note that no binarization is available for SparCC, *i*.*e*., is a weighted network. The same pre-processing steps of MAGMA network, *i*.*e*., prevalence and abundance filters, are applied to SparCC microbial network. We notice that intra-phylum connections are stronger than inter-phyla. SparCC connections are, on average, stronger than in the weighted MAGMA’s counterparts. As for MAGMA, *Proteobacteria* and *Bacteroides* yield stronger intraphylum connectivity than *Firmicutes*. Only one microbe (lowest annotation: Class *TM7-3*) belongs to the *TM7* phylum, and shows increased connections with *Firmicutes, Proteobacteria*, and *Verrucomicrobia* phyla at 9m. Moreover, it is interesting to note that for both MAGMA and SparCC networks, the connections involving *Proteobacteria*’s taxa are stronger at timepoint 9m.

### 3.3 Longitudinal Analysis: Neighbourhood Dynamics in Global Networks

We applied MNDA to a multiplex network with two layers, each consisting of the global microbiome co-occurrence networks at month 6 and month 9, and computed cosine distances between all possible pairs of microbial taxa in the embedding space (Section 3.2). The measures of dissimilarity can be used to cluster taxa pairs. Pairs may involve components from the same layer or different layers of the network. However, neural networks usually have a non-convex cost function; thus, reruns of the EDNN can lead to different outcomes. Therefore, multiple repeats of MNDA were used as inputs to a novel implementation of robust clustering (see next – *a novel implementation of ensemble consensus clustering*). This led to two robust clusters, shown in Figure 7A as *cluster 1* and *cluster 2*. The confusion table capturing how many matched months 6-9 pairs of microbial taxa belong to the same cluster or are spread over two clusters, is given in Figure 7B. Corresponding transition probabilities (*P*_*i*_ → *j* = *P* (cluster *j* at 9*m* | cluster *i* at 6*m*)) are visualized in Figure 7C. These show that the majority of microbes have a similar global network neighbourhood dynamic. For 24 out of 95 microbial taxa this is not the case. These are listed in Additional file 3: Table SI, with their corresponding genus-species names. The two microbial taxa with similar global neighbourhoods, yet different from those microbes in cluster 1, are *Bacteroides uniformis* and *Blautia sp*. We identified microbes with high or low neighbourhood dynamics by sorting the robust co-clustering similarities for each time-matched pair of taxa. In particular, we selected the first larger jumps at both extreme ends of the similarities, respectively, as shown in Additional file 3: Figure S8A. A complete list of microbes with extreme neighbourhood dynamics is given in Figure 5D and Table I, which also lists *appearing* and *disappearing* microbes (defined in Section 2.2 – *pre-processing*). Notably, rather than using co-clustering similarities to rank taxa in terms of their dynamics, we could also have ranked taxa directly via their corresponding cosine distances in the MNDA embedding space, averaged across multiple MNDA runs, and re-ranked. This procedure led to similar results (not shown).

**Table 1.**
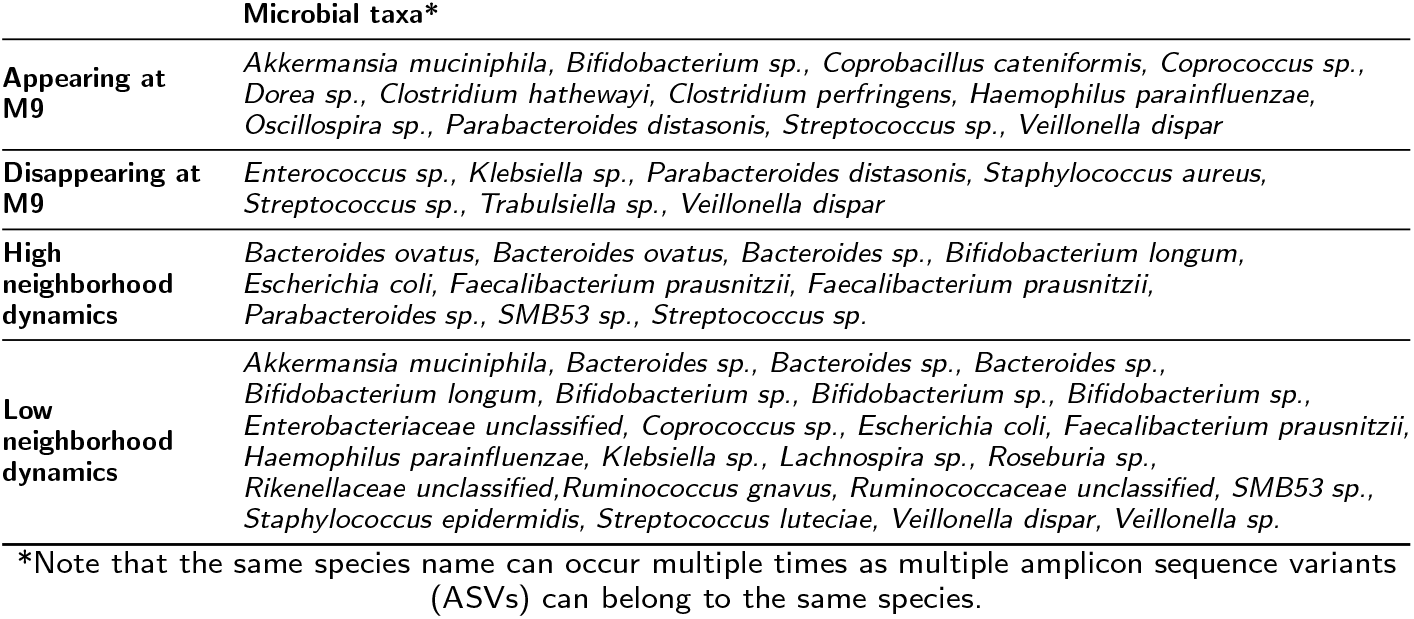
The list of highly dynamical microbes with their corresponding genus-species names.

|  | Microbial taxa* |
| --- | --- |
| **Appearing at M9** | *Akkermansia muciniphila*, *Bifidobacterium* sp., *Coprobacillus cateniformis*, *Coprococcus* sp., *Dorea* sp., *Clostridium hathewayi*, *Clostridium perfringens*, *Haemophilus parainfluenzae*, *Oscillospira* sp., *Parabacteroides distasonis*, *Streptococcus* sp., *Veillonella dispar* |
| **Disappearing at M9** | *Enterococcus* sp., *Klebsiella* sp., *Parabacteroides distasonis*, *Staphylococcus aureus*, *Streptococcus* sp., *Trabulsiella* sp., *Veillonella dispar* |
| **High neighborhood dynamics** | *Bacteroides ovatus*, *Bacteroides ovatus*, *Bacteroides* sp., *Bifidobacterium longum*, *Escherichia coli*, *Faecalibacterium prausnitzii*, *Faecalibacterium prausnitzii*, *Parabacteroides* sp., *SMB53* sp., *Streptococcus* sp. |
| **Low neighborhood dynamics** | *Akkermansia muciniphila*, *Bacteroides* sp., *Bacteroides* sp., *Bacteroides* sp., *Bifidobacterium longum*, *Bifidobacterium* sp., *Bifidobacterium* sp., *Bifidobacterium* sp., *Enterobacteriaceae unclassified*, *Coprococcus* sp., *Escherichia coli*, *Faecalibacterium prausnitzii*, *Haemophilus parainfluenzae*, *Klebsiella* sp., *Lachnospira* sp., *Roseburia* sp., *Rikenellaceae unclassified*, *Ruminococcus gnavus*, *Ruminococcaceae unclassified*, *SMB53* sp., *Staphylococcus epidermidis*, *Streptococcus luteciae*, *Veillonella dispar*, *Veillonella* sp. |

\*Note that the same species name can occur multiple times as multiple amplicon sequence variants (ASVs) can belong to the same species.

**Figure 7.**
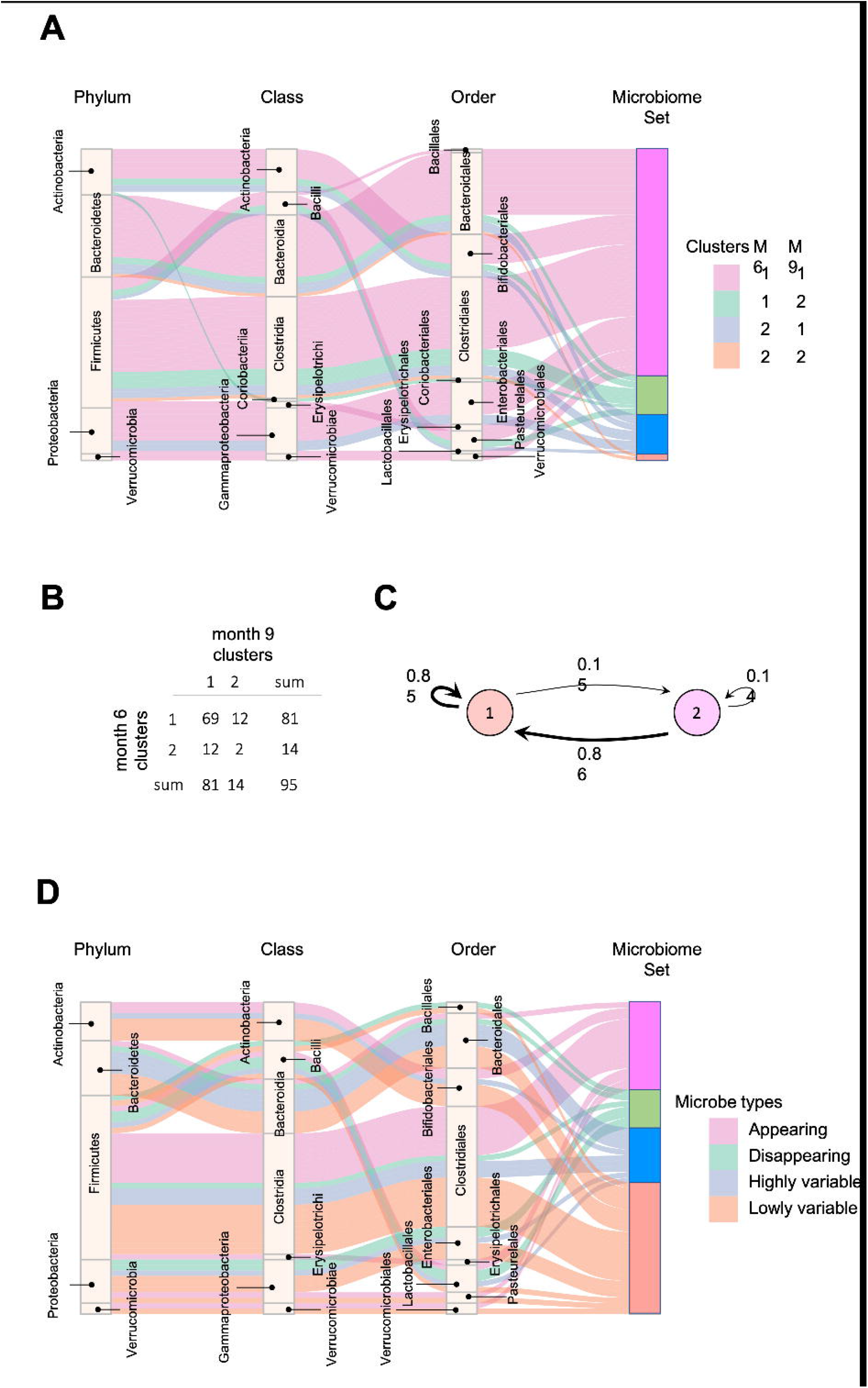
(A) Four groups of microbes based on their cluster shifts between m6 and m9; (B) The frequency of cluster shifts between m6 and m 9; (C) Graphical illustration of microbial cluster shift rates; (D) Highly and lowly variable microbes.

Overall, *Firmicutes*’ microbes constitute the vast majority of appearing (69%) and disappearing (57%) microbes, *i*.*e*., microbes that would pass the 15% prevalence threshold only at month 9 (appearing), or at month 6 (disappearing). Microbes with high (low) neighbourhood dynamics constitute 11% (25%) of 95 original microbes. Phyla *Actinobacteria, Bacteroidetes, Firmicutes, Proteobacteria*, and *Verrucomicrobia* deliver respectively 1.6% (6.6%), 1.8% (1.8%), 0.6% (1.6%), 0.7% (2.8%), and 0.0% (25%) of high (low) dynamic microbes in terms of their global network neighbourhoods.

#### A novel implementation of ensemble consensus clustering

Many clustering methods can be made more robust by *consensus clustering* (CC). Here, we propose a CC framework to achieve robustness by repeatedly applying MNDA (50 repetitions) and subsequent clustering of microbes based on their cosine distance. As mentioned before, cosine distance is computed across all pairs of possible microbes, spanning data on months 6 and 9. The adopted clustering strategy is *k-means*; the number of clusters *k* is determined by maximising the *Silhouette index*. For M microbes, of which dynamics need to be tracked across time points, we calculate the 2*M ×* 2*M* matrix of co-clustering. Matrix entries are the frequencies a pair of microbes (a single microbe at two time points or two microbes at some point in time) jointly belong to the same cluster among all clusterings. This co-clustering matrix can be viewed as a robust similarity matrix on the basis of which we can define a robust “distance” between any pair of nodes in the original MNDA embedding space. The more distant two nodes in the embedding space are, the more their local neighbourhoods differ. Hence, when two nodes refer to the same microbe, measured at different time points, high co-cluster similarity implies low local neighbourhood dynamics for that particular microbial taxon, considering its global microbiome co-occurrence neighbourhoods at months 6 and 9. We implemented consensus clustering proposed by Monti et. al [46] on the co-clustering matrix to obtain the optimum number of clusters along with the clustering results. As depicted in Additional file 3: Figure S8B, the algorithm suggests two clusters of microbes by calculating the area under the curve of the *empirical cumulative distribution*. For additional details we refer to Monti et al. [46].

### 3.4 Longitudinal Analysis: Neighbourhood Dynamics in Individual-Specific Microbiome Networks

An alternative view on microbiome data is given by ISNs, which provide edge information at an individual-specific level. Hence, where in Section 3.4, time-course analyses involved comparing global microbiome co-occurrence networks at each time point, here, we do so at an individual-specific level (Figure 8A). We used the same MNDA framework, but instead of submitting a single multiplex network (Section 3.3), we submited 69 multiplex ISNs simultaneously. We show how our new notion of microbiome neighbourhood dynamics (explained in Section 2.5), when assessed on a per-individual level, may offer complementary views compared to standard data views. The most standard data view analyses (transformed) microbial abundances for study subjects; data is organized according to a matrix depicted in (Figure 8B), where the microbial abundances of both timepoints are considered as features (node-oriented approach). Since the rise of individual-specific network construction techniques (outlined in the Background Section), data records may additionally (or only) involve information about individual-specific edges (presence or absence, or edge strength on an interval scale). Such an edge-oriented data format is depicted in (Figure 8C), where the features are the microbial co-occurrences of both timepoints (edge-oriented approach). With the newly proposed MNDA framework, dynamics across time points is investigated in an embedding space and each individual can be assigned a vector of cosine distances. As features, each cosine distance captures ISN local neighbourhood dynamics across time points (Figure 8D – dynamic-oriented approach). The similar data formats between Figure 8B, 8C, and 8D allow adopting similar association modelling or prediction modelling strategies, yet interpretations will differ. It is noteworthy that our proposed method results in the least feature size as we have one feature for each taxon independent of the number of time points.

MNDA-induced prediction outperforms other methods and complements standard approaches. To support this statement, we developed prediction models for mode of delivery (C-section versus vaginal) and diet type (persistent versus non-persistent – as defined in Section 2.1). In particular, we applied *support vector machines* (SVM) with *radial basis function* (RBF) kernels to the data organized in each of the aforesaid structures (Figure 8). In the training phase, we balanced the classes’ size via under-sampling of the majority class. To reduce the dimensionality of the data, we used a *forward feature selection* framework. We repeated the modelling process, each time leaving out a single individual, as part of a leave-one-out cross-validation. The left-out individual was used to test the trained model. The entire process was repeated 100 times; AUCs were averaged across repeats and standard errors computed. The results are reported in Figure 9. MNDA-informed prediction models consistently outperform models that only either use microbial abundance or MAGMA individual-specific edge weights as input features. The advantage of using individual-specific edges is context-dependent: depending on the timepoint and the phenotype, the classification performances vary. The edges at m9 have the best performance among MAGMA individual-specific edge weights with AUC of 0.57 and 0.64 for mode of delivery and diet. However, it is underperforming compared to directly using CLR-transformed microbial abundances at m9.

**Figure 8.**
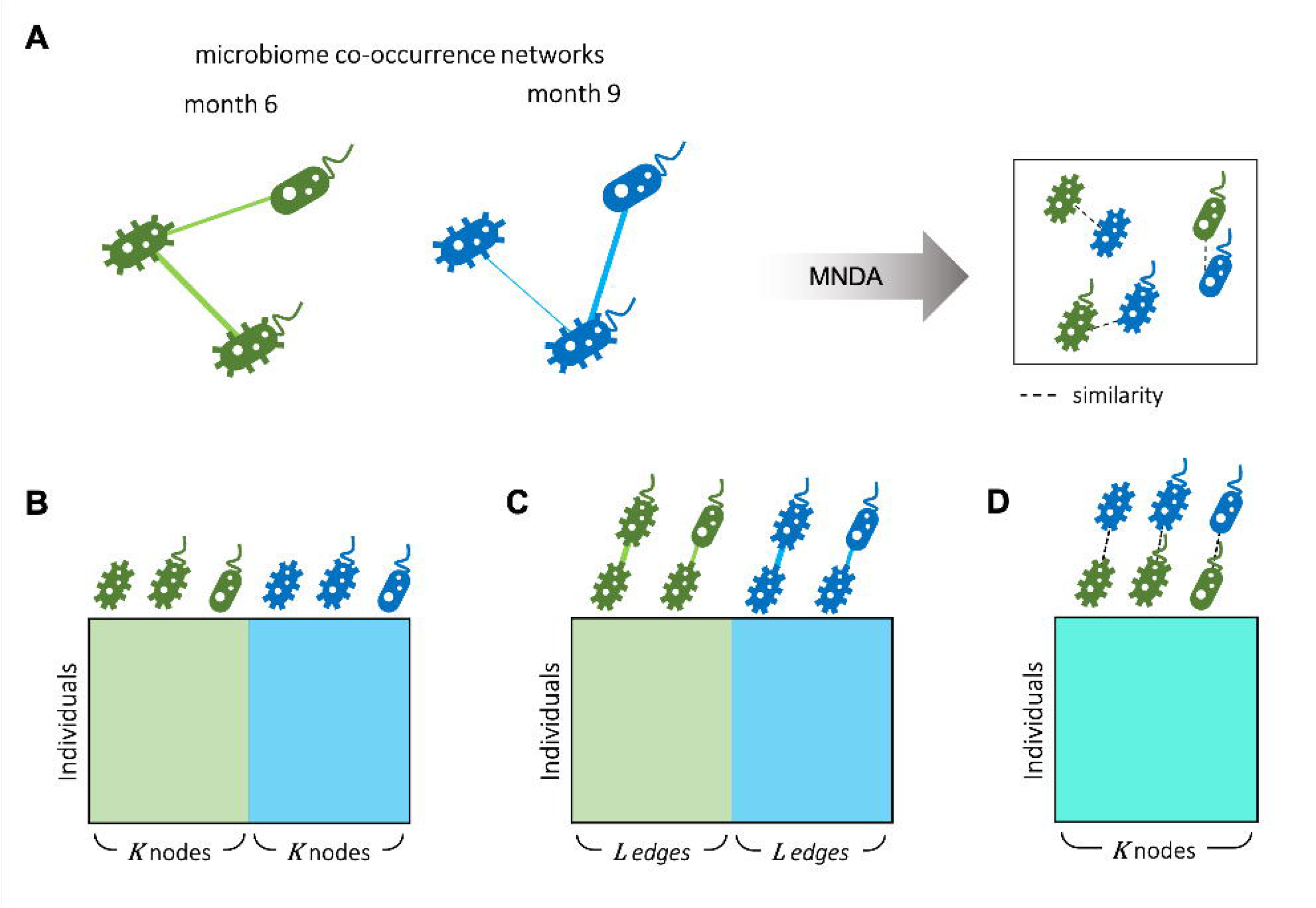
Different scenarios for microbiome longitudinal analysis. (A) Individual specific networks of microbiome co-occurrences for month 6 and month 9 are represented into an embedding space using our proposed MNDA method. Therefore, data can be organised by different perspectives as following. B) node-oriented approach: standard Taxon abundance table of both timepoints is used; (C) edge-oriented approach: the edge weights of both timepoints are used as features; (D) dynamic-oriented approach: the variations between the local neighbourhood of nodes in time are considered as features. Assuming K nodes and L edges between nodes, the number of variables in the microbial dynamic space is the same as the number of microbes K. The number of edges L is bounded by the number of possible selections of pairs of nodes out of K nodes.

**Figure 9.**
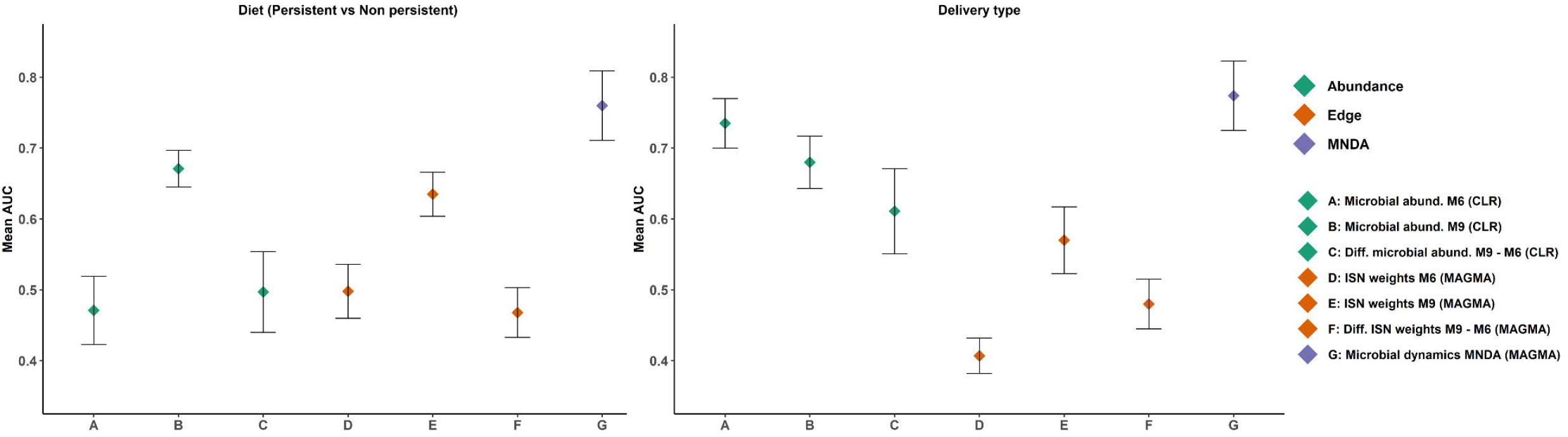
The prediction results, in terms of AUC, for (A) delivery type and (B) diet type. Different feature sets use for prediction, among which MNDA-based method is consistently the top performing. On the contrary, the difference between MAGMA individual-specific edges at 6m and 9m never reaches an AUC of 0.5.

ISN dynamic analysis highlights microbes not identified via global network analyses. Important discriminative microbes for diet type or mode of delivery were identified by counting the number of times a microbe was selected by the adopted forward feature selection algorithm, mentioned above, out of 69 runs (every run had one individual being considered as validation sample) and 100 repeats. It generates a ranking of microbial importance: the higher the selection count, the higher the microbe’s importance. Results over 50 generated embedding spaces were summarized via summing 345,000 repeats per microbe, giving rise to a robust final ranking of important discriminative microbes. We emphasize that the selection of microbes in discriminative models was based on a measure of local neighbourhood dynamics across time points. A list of top discriminators in this sense is provided in Table II, using annotations of genus-species names. Among these taxa, *Lachnospira sp*. and *Bifidobacterium sp*. have low neighborhood dynamics; besides, *Streptococcus luteciae* and *Ruminococcus sp*. are important microbes for both delivery and diet types. The latter neighbourhood dynamics analysis did not account for differences between infants by diet during months 6 and 9, nor delivery mode. Specifically, *Streptococcus luteciae* had previously been reported to be associated with infant feeding [47]; moreover, its association with the delivery type can be explained by its relation to the skin bacterium.

**Table 2.** The list of top ranked discriminator microbes (genus-species names) for delivery type and diet mode.

|  | Microbe ASVs |
| --- | --- |
| **Delivery type** | *Streptococcus luteciae*, *Trabulsiella* sp., *Ruminococcus* sp., *Ruminococcus* sp., *Parabacteroides distasonis*, *Lachnospira* sp., *Bifidobacterium* sp. |
| **Diet Mode** | *Bacteroides ovatus*, *Clostridium citroniae*, *Bacteroides ovatus*, *Bifidobacterium bifidum*, *Ruminococcus* sp., *Streptococcus luteciae*, unclassified *Lachnospiraceae*. |

Stratified analyses confirm the differential dynamic behaviour of identified discriminators in Table II. When a microbe is highly discriminative for diet type and discrimination is based on ISN local neighbourhood dynamics over time, then the change over time of its immediate “interaction” partners should also be markedly different between dietary strata; and the same for mode of delivery. In Figure 10, we focus on the microbial taxa of Table II (dark nodes) and their distance-1 neighbours to investigate how the edge weights change over time. For each class of delivery mode (C-section or vaginal delivery) and diet mode (persistent or non-persistent) we obtained two subnetworks for each time point, month 6 and month 9, by averaging ISNs from infants belonging to the same class per time point. Then, we subtract the edge weights of networks in month 6 from the edge weights of networks in month 9, resulting in four networks for each class. In Figure 10, the presented edges strengths indicate the differences between averaged edge strengths between time points for each class (red for average edge weight at 9m larger than at 6m and green for the reverse). Each node is annotated with its genus-species name. For further comparison of these difference networks within the delivery mode and diet, we refer to Additional file 3: Figure S9. We observe that the differences in co-expression networks over time show that the connections at 6M are much stronger than at 9M for C-section delivery. Furthermore, we observe that a change in diet between the two timepoints results in clear differences in co-expression networks (Figure 10C), while the networks are apparently more stable when infants have a more stable dietary pattern.

**Figure 10.**
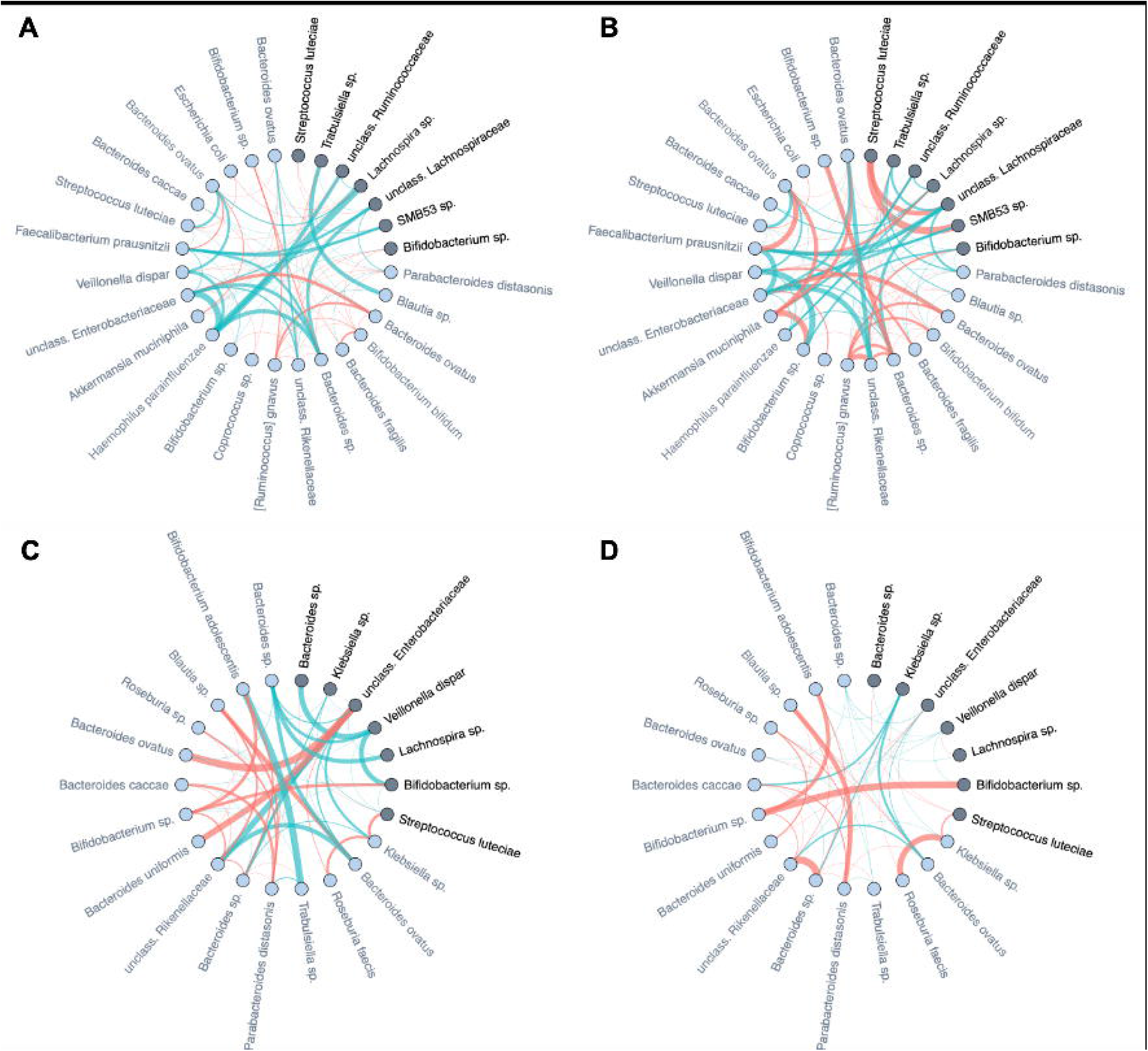
Differences of averaged microbial co-expression networks restricted to the important microbes and their first level neighbours between month 6 and month 9 on (A) C-section and (B) vaginal delivery, as well as (C) non-persistent diet and (D) persistent diet. Edge thickness is given by its co-occurrence magnitude, while the edge colours show the sign of the correlation (red for average edge weight at 9m larger than at 6m and green the reverse). The important microbes are highlighted by dark colour and their family names (also listed in Table II), and their first level neighbours are indicated by light colours.

Specifically for the diet mode, intra-class variation of ISNs restricted to the same considered taxa in Figure 10 is illustrated via *so-called graph filtration curves* (Figure 11) [48]. These curves provide a more refined or complementary view to the averaged ISN representations in Figure 10: Each individual’s contribution to the average can be shown. These filtration curves show that the largest variation between the two timepoints is observed for non-persistent diet (*i*.*e*. the variation in the diet mode is correlated with the dynamics of the microbial co-occurrence). Hence, our analysis confirms that dietary shifts have a stronger impact on dynamics of local neighbourhood microbial co-occurnences than observed between groups characterized by different modes of delivery.

**Figure 11.**
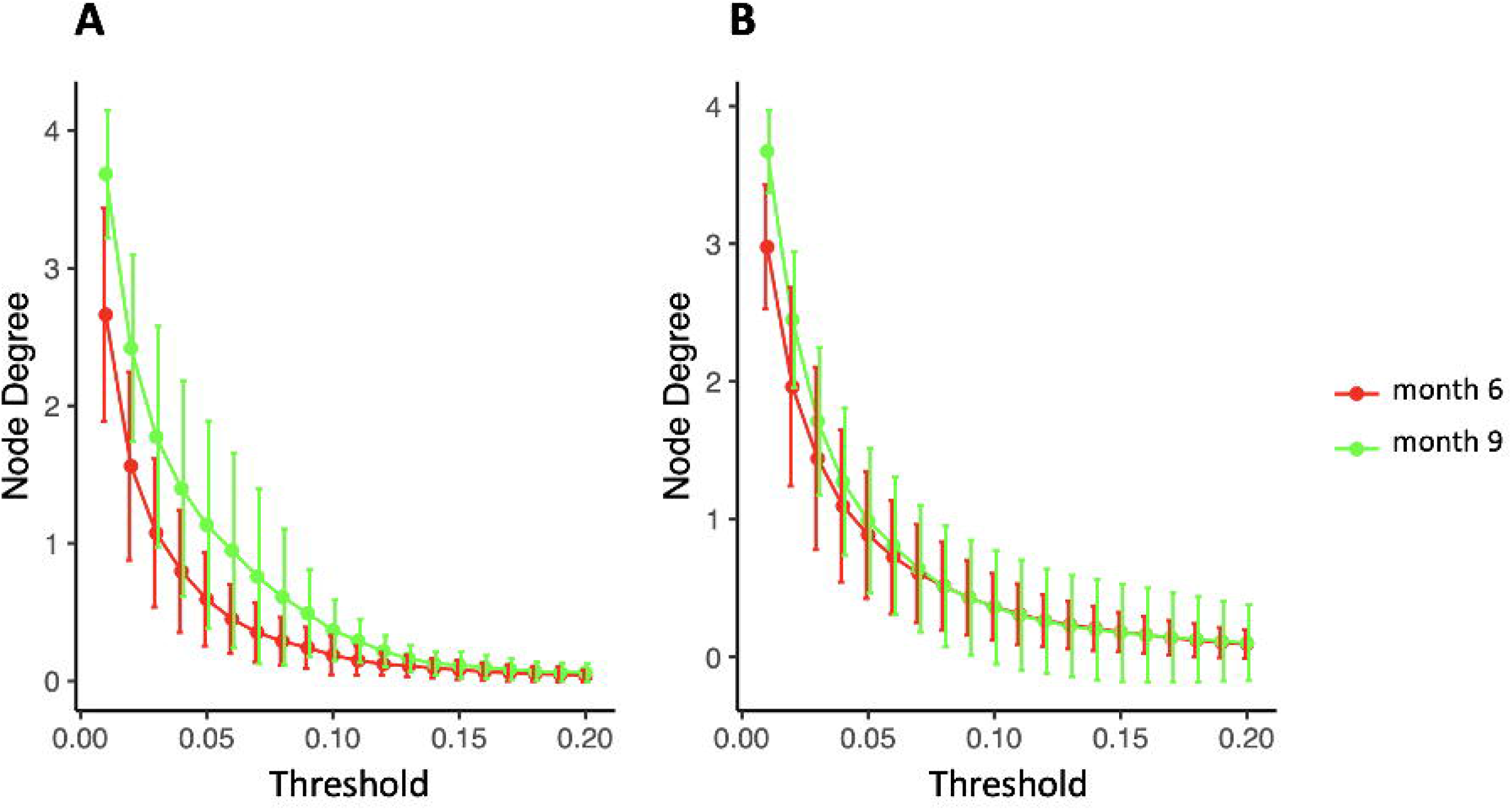
Filtration curves representing the difference between averaged microbial co-expression networks of month 6 and month 9 for (A) non-persistent diet (9 infants) and (B) persistent diet (32 on solid food, 27 on mixed breastfeeding and solid food).

### 3.5 Local Neighbourhood Dynamics to Identify Between-Individual Heterogeneity

MNDA-based similarity measures can be used to cluster individuals into homogeneous groups according to similar microbial neighbourhood dynamics. Using ISNs to define neighbourhoods, we observed that clusters of individuals significantly differed from those obtained via *Dirichlet multinomial mixtures methodology* (DMM) [32]. More specifically, DMM clustering on pooled data across time points revealed two clusters. These are depicted in Figure 12, together with corresponding transition information as infants grew older. In contrast to DMM, MNDA-induced clustering groups individuals according to their dynamics-similarity in microbial interaction patterns. Robust clustering was performed, as described in Section 3.3 – *a novel implementation of ensemble consensus clustering*. This analysis also highlightes two clusters, roughly of the same size (33 and 36 individuals). We used a Chi-square statistic to evaluate the degree of non-random correspondence between MNDA-or DMM-clusterings. This gave rise to a permutation-based *p-value* of 0.2814. It is worth noting that, unlike DMM, MNDA results in communities that do not change over time; they are based on dynamic information across time points. Microbial abundance changes over time and dynamics of microbial interactions are two distinct reflections of the same process.

**Figure 12.**
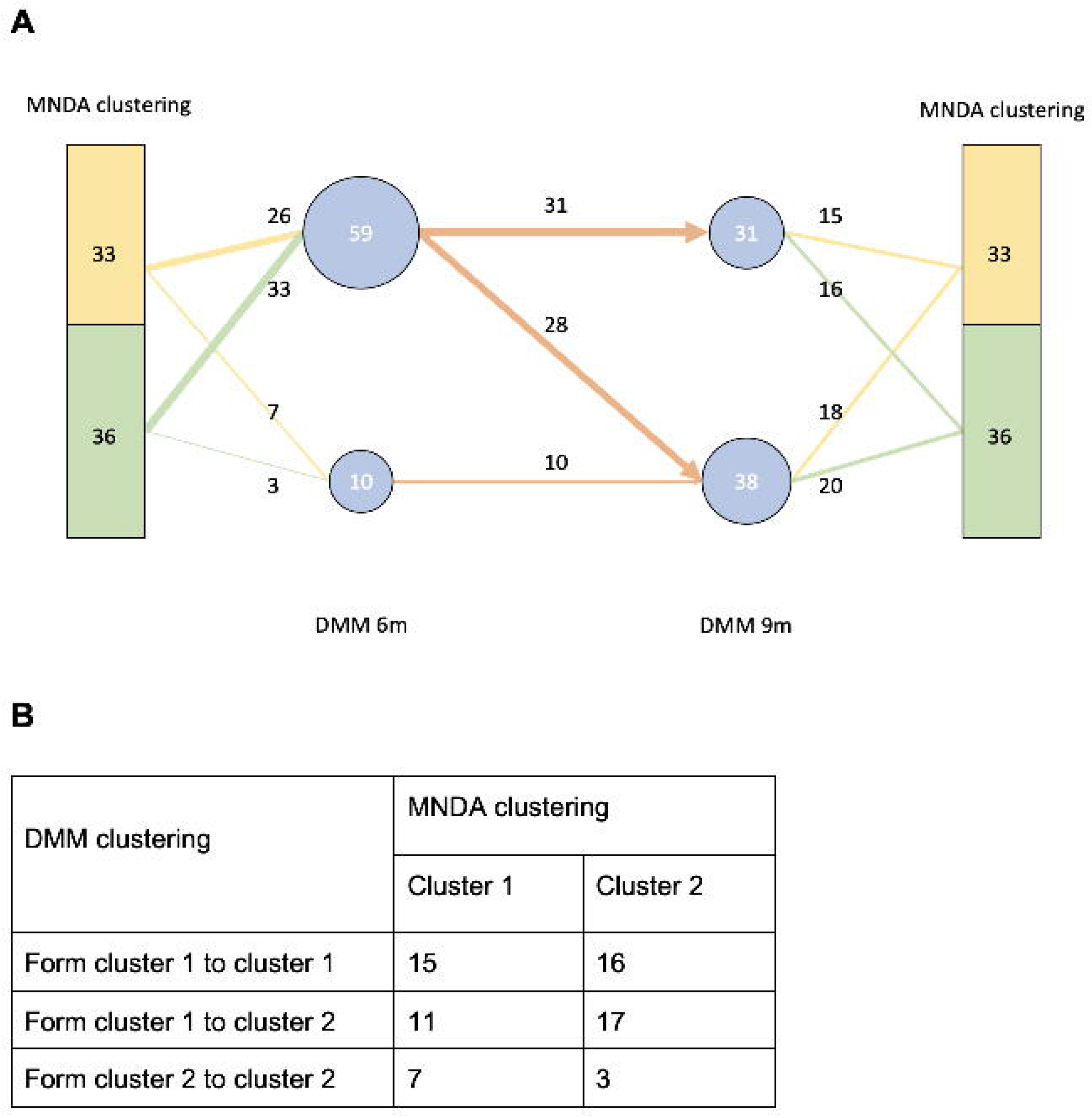
A comparison between our MNDA-induced clustering and DMM clustering reveals no significant association, which means that MNDA provides an independent view to the data. It is also noteworthy that unlike DMM that finds clusters for each timepoint, MNDA provides clustering of individuals based on their variation in both timepoints.

## 4 Discussion

### 4.1 The Value of Individual-Specific Dynamic Microbial Networks

Environmental factors may play a critical role in human complex diseases [49, 50]. One of these critical factors, the gut microbiome, has received special attention in recent years, for instance in the context of disease development and progression [51, 52]. Even though many studies have shown the resilience of the gut microbiota and its stability over time, the gut microbiota is subject to dramatic shifts due to person-oriented interventions such as changes in diet or medication use.

In this work, we proposed a novel work analysis framework, referred to as MNDA, to capture the dynamics of gut microbial co-occurrence, within and across individuals. The approach is built on the concept of individual-specific microbiome networks, with microbial taxa as nodes and individual-specific connecting edges defined via microbial interactions. Microbiome ISNs were embedded in a common space. The cosine distance of the same node (microbe) between two timepoints in the embedding space was taken to quantify local node-neighbourhood dynamics. This information was subsequently exploited to stratify individuals into different homogeneous subpopulations, revealing new aspects of population heterogeneity from the microbiome interactome perspective. The proposed strategy was illustrated on data from the LucKi cohort [33], containing microbiome profiles of 69 newborns collected at two different time points (6 and 9 months after birth). Via comparison with baseline techniques in the field, we motivated the potential of microbiome ISNs in microbiome time-course analyses.

Numerous studies have been performed on microbial longitudinal taxon abundance data, amongst others, in association with clinical outcomes. However, these studies typically ignore microbial interactions, the dynamics of which could also be highly informative. Taking such microbial co-occurrences into account when modelling temporal dynamics in bacterial communities, *generalised Lotka-Volterra* models [53, 54] and *dynamic Bayesian network* models [17, 55] have been developed. Even though these methods may describe the development of dynamic microbial networks, data are aligned by assuming that patterns are similar across individuals, yet exhibit different rates of change in accordance with demographic and clinical variables. Individual-specific microbe neighbourhoods in ISNs are ignored. Also, these methods typically require a large number of timepoints, which may not always be available in microbial cohorts dealing with humans.

Our MNDA framework was exemplified on time course data with two time points. We selected cosine distance as a measure of microbial neighbourhood dynamics across two time points, as we aimed to capture the amount of similarity between two data points in a two-dimensional subspace of the MNDA embedding space. Cosine distance is linked to, but not the same as, angular distance, which varies in the range [0, 1]. Angular distance is a true metric but requires the computation of *arccos*(.). When cosine distance between data points in the MNDA embedding space is small (or equivalently, cosine similarity is high), then the two points will be located in the same general direction from the origin. In other words, one can be seen as a scaled-up version of the other.

### 4.2 The Value of Advanced Representation Learning

At the heart of MNDA lies an encoder-decoder algorithm to embed local microbial neighbourhoods at multiple timepoints. Neighbourhoods may be based on global microbiome co-occurrence networks or on derived ISNs. Traditional embedding approaches [56], such as PCA of Laplacian matrix, *DeepWalk* [57] and *node2vec* [58], have been previously proposed for single-layer networks. For these methods, the parameters in the encoder are not shared between nodes and cannot be generalised to new nodes. Using an EDNN structure, MNDA is able to introduce shared parameters while encoding the nodes. In addition, we solved the node generalisation problem by feeding the vector of node neighbours to the encoder. Using this trick, the encoder can generate embedding for a new node whose neighbours need to be determined. In addition, the method can easily be extended to multiple timepoints by increasing the number of layers of the multiplex network. Our MNDA implementation works with weighted networks by exploiting a fixed-length weighted random walk algorithm. Even though MNDA can be used on binarized networks as well, with binary input and binary output vectors for the encoder-decoder system [59], we do not describe it here since our empirical evaluation indicate suboptimal results.

Computing ISNs can be computationally intensive when the number of edges grows. On a computational infrastructure with *Windows 10* and *R version 4*.*0*.*3 (2020-10-10)*, computing 5 ISNs for 95 microbes took 1.17 seconds. The MAGMA network calculation can rapidly become intractable: for 5 MAGMA networks on 95 microbes, it took 55.15 seconds. MNDA analysis, including random walk probability calculation and EDNN training, is the most time-consuming step in our analysis. In particular, on a *MacOS (version 12*.*4)* and *R version 4*.*0*.*5 (2021-03-31)*, random walk probability calculation took 28.1 seconds for each multilayer network of 95 microbes; each round of EDNN training took 5.7 minutes for 69 individuals. The measurements were done through the *Sys*.*time()* function in *R*. Embedding individuals into a joint space is less computationally intensive than creating an embedding space for each individual separately. It also ensures that cosine distances are comparable across individuals (and thus can be used to cluster individuals). However, on the downside of this, when the initial dataset is enriched with additional individuals, the joint embedding space will differ, and individual predictions may change.

### 4.3 Unique Use of Individual-Specific Network Methodology

The ISNs were constructed using the method by Kuijjer et al. [20]. It builds on a global reference network from which individuals are iteratively extracted and perturbation effects are used to derive an ISN with individual-specific edges. Motivation for this particular way of ISN construction included that it is easy to implement via the LIONESS software [21] and has a straightforward interpretation: On average, and for an asymptotically large number of individuals, Kuijjer’s ISNs average to the global reference network. Moreover, unlike for sample-specific networks, approaches such as described in [60], edges in Kuijjer’s ISNs go beyond differential co-occurrences: for each individual a tailored “co-occurrence” network is reconstructed.

Given the compositional and zero-inflated nature of the microbial data, it is necessary to use a global network association algorithm that accommodates these data characteristics. One of the most common tailored microbial association network inference approaches in the field is SparCC [61, 62]. Alternatively, Cougoul et al. [39] proposed MAGMA (*rMAGMA R* package), which not only takes into account microbiomes’ noisy structure, excess of zero counts, overdispersion (high skewness) and compositional nature (simplex constraint), but also ensures inferred association strengths within [−1, 1]. Moreover, it results in a sparse low dimensional matrix of co-occurrences, with edge strengths that can be adjusted for known confounders. On the contrary, SparCC does not have a native sparsification and needs to be added as an external step, with no established rule on how to perform it, hence, undermining the reproducibility. Since ISNs are computed outside the MNDA framework, our pipeline can be combined with any type of ISN, as long as the edges are individual-specific.

Compared to the work done by Mac Aog’
sain et al. [24], our work has some important differences. First, the authors construct microbiome co-occurrence networks on CLR-transformed abundances and Pearson correlation. Pearson correlation is the base measure of association implemented in the LIONESS software; which was initially showcased on gene expression data. Friedman et al. [61] showed that even though correlations on the CLR transformation are more accurate than Pearson correlation, they are not as accurate as the SparCC algorithm. Second, the authors continued their analysis with edges as units of analysis. Our work exploits individual-specific networks to define individual-specific ASV neighbourhoods (hence sets of edges). Third, MNDA is mainly developed to compare microbiomes across conditions or over time, by stacking ISNs into multiplex networks. Compared to the iENA protocol [26] applied to multi-time point microbial analysis [27], our newly proposed workflow is different in the following ways: *i)* firstly, the MAGMA transformation is considered instead of a Pearson correlation on abundances; *ii)* LIONESS aims to reconstruct a network with the same interpretation as the global network, *i*.*e*., the population-based network, while iENA computes an individual deviation from the average; *iii)* edge-network in iENA is aggregated in a single sCI value quantifying the disease’s risk, while in our work the edges constitute the input of MNDA pipeline. They state that only interactions, not abundances are significant and suggest remarkable disruption of the microbiota community when diseases occur. This understanding is reinforced by a follow-up work from the same group, characterising the personalised microbiota dynamics for disease classification by individual-specific edge-network analysis [25]. Beneficial co-evolved interactions between host and microbiota can be disrupted by different environmental stresses such as changes in dietary habits, natural physiology, virus infections, and medical treatments [63, 64, 65]. In general, analysis techniques developed to process timevarying networks often require numerous temporal observations. The analysis of time-dependent multiplex networks with low temporal dimensions remains largely under-investigated. In this paper, we encapsulate our proposed time-course analysis of ISNs into a multiplex network differential analysis framework, where each network layer refers to a point in time. The framework quantifies the changes in the local neighbourhood of each node for an ISN (*i*.*e*. a microbial taxon) between the time points. This is achieved by embedding the nodes of network layers into a shared embedding space.

### 4.4 Complementary Novel Findings

We uncovered previously unreported microbial taxa as biomarkers for temporal changes between months 6 and 9 after birth. In particular, the two microbial taxa that consistently cluster together in months 6 and 9 and are different from other microbes are *Bacteroides uniformis* and *Blautia sp*. We can find insights from the composition of the appearing/disappearing microbes. The disappearing *Firmicutes* microbes’ are mostly (*facultative*) aerobic microorganisms, including *Enterococcus, Streptococcus*, and *Staphylococcus* species, along with other *facultative aerobes* such as *Klebsiella* (phylum *Proteobacteria*). In contrast, the appearing taxa within the *Firmicutes* phylum include several strictly anaerobic species, including *clostridial* members, *Dorea sp*. and *Coprococcus*. This shift is indicative of a more reduced intestinal environment and more mature *microbiome* adapting to the fermentation of complex dietary carbohydrates. Moreover, *Verrucomicrobia* is recognized as the phylum with the least dynamic microbes in terms of their global network neighbourhoods.

Regarding the individual-specific neighbourhood dynamics, *Lachnospira sp*. and *Bifidobacterium sp*. have low neighborhood dynamics; *Streptococcus luteciae* and *Ruminococcus sp*. are important microbes for both delivery and diet types. *Streptococcus luteciae* had previously been reported to be associated with infant feeding [47]; and, it is a skin bacterium, which can be the reason why in our study it was associated with birth mode (like other skin bacteria) and exhibited low neighbourhood dynamics. Other markers, such as *Ruminococcus sp, Lachnospira* and *Bifidobacterium sp*., are too general, and little biological interpretation can be extracted in the context of this study. However, the observed link of *S. luteciae* and *B. bifidum* with diet can potentially trace back to the depletion of these microbes once breastfeeding is ceased.

The differences in co-expression networks over time (as highlighted in Figure 10) show that the connections at 6M are much stronger than at 9M for C-section delivery. This is indicative of the waning effect of C-section delivery and temporal colonization of environmental and skin bacteria in C-section delivered infants that are displaced by other bacteria. The interaction networks of such typical C-section delivered microbes also appear to wane over time. This is exemplified by the edge between *Lachnospira sp*. and *Haemophilus parainfluenza*, a bacterium typically found to be temporarily enriched in C-section infants, which is much stronger at 6 as compared to 9 months of age. Furthermore, we observed that a change in diet between the two timepoints results in clear differences in co-expression networks (Figure 10C), while the networks are apparently more stable when infants have a more stable dietary pattern. Note that even infants with persistent dietary patterns still have a diet that is gradually becoming more complex and diverse as more complementary foods are being introduced over time.

Moreover, filtration curves indicate that newborns shifting diet between 6M and 9M have the largest variation between the two timepoints. This would indicate that dietary shifts have a stronger impact on dynamics of microbial co-occurrence in this age-window when compared to C-section, which is in line with previous studies indicating that the impact of birth mode is mainly restricted to the first months of life.

### 4.5 Limitations and Future Work

Several steps in our MNDA framework can be varied. We could have chosen Euclidean distance instead of cosine distance. However, Euclidean distance calculations are computationally intensive and are often replaced by Manhattan distance for high-dimensional data. In scenarios of high dimensionality, the approximation error introduced by Manhattan distance may be unacceptably large and thus undesirable. For the user who does wish to adopt Euclidean distances instead, Carderilli et al [66] proposed an approximation method to Euclidean distance in high-dimensional spaces. Building upon our default implementation of cosine similarity, in principle, we can generalize the adopted similarity measure to more than two time points by moving from angular similarity to similarity based on *dihedral angles* (*i*.*e*., angle between two intersection half-planes) or generalized solid angles of pointed convex cones (*i*.*e*., the intersection of a finite number of half-spaces whose corresponding hyperplanes meet in exactly one point [67]). Assessing performance of multiple distance measures to capture microbiome neighbourhood dynamics as a biomarker for prediction or stratified medicine is a logical next step.

Microbiome data analysis results are particularly susceptible to choices made during virtually all steps of the analysis: for instance during pre-processing (*e*.*g*., normalization [68]), when adopting differential abundance strategies [69], or when carrying out network analyses [29]. ISNs can be computed in several ways, with individual-specific edges being binary or weighted, sparse or rich, positively weighted or not. In this work, we transformed ISN edge weights to their absolute values. Hence, we did not differentiate between positive and negative correlations. Loftus et al. [70] noticed that taxonomically and functionally similar species tend to have positive associations. In the current version of MNDA, taxa with neighbourhoods at months 6 or 9 only differing in sign, would be considered highly similar in terms of their neighbourhood dynamics. In future work, we aim to adapt MNDA to accommodate positive and negative edge weights. Particularly linked to our MNDA framework, tuning the hyperparameters of EDNN may further enhance performance. These hyperparameters are number of hidden units (dimension of the embedding space), number of layers, *L*_1_ and *L*_2_ regularization parameters, and batch size. Although performance can be clearly defined in view of expected prediction accuracy in supervised context (see Section 3.4 – *MNDA inspired prediction*), it is less clear in unsupervised modelling contexts, in the absence of the ground truth. For instance, the relevance of homogeneous subgroups identified in Section 3.5 may become more apparent when associated with variation in extraneous data. For this study, we only had additional information about diet and mode of delivery. No significant association was observed between diet, mode of delivery and cluster membership (using chi-squared test *α* = 0.05).

Finally, we emphasize that our MNDA framework is not limited to temporal microbiome data. For instance, nodes can be genes with edge weights representing gene co-expression. ISNs can represent molecular interactions within and between multiple related omics data types [71]. For all these scenarios, we believe that our longitudinal analysis framework can be useful to identify novel biomarkers for precision medicine.

## 5 Conclusion

In this paper, we propose a novel framework to uncover microbial neighbourhood dynamics. Our approach, MNDA, combines representation learning and individual-specific microbiome networks, which makes it unique in the current landscape of statistical methods for microbiome temporal data analysis. MNDA is not restricted to microbiome data but can handle any data type and measurements, as long as these can sensibly be organized into cross-sectional association networks. Our results show that MNDA can induce predictions that outperform standard approaches and that ISN dynamic analysis can identify microbes that are not identified by global network comparisons. Stratified analysis over clinical variables confirms the differential dynamic behaviour of identified discriminators to diet type stability or mode of delivery. Standard microbial abundance changes over time and MNDA dynamics of microbial interactions can be seen as alternate representations of the same underlying process.

## Ethics approval and Consent to participate

The LucKi Birth Cohort Study was approved by the Medical Ethical Committee of Maastricht University Medical Centre (MEC 09–4–058). LucKi is designed according to the privacy rules that are stipulated in the Dutch ‘Code of Conduct for Health Research’ [33].

## Consent for publication

Not applicable.

## Availability of data and materials

The global network figures are generated by the *rMAGMA* graphical tool and the *igraph R* package. The averaged co-expression networks are generated by the *R* package *ggraph*. The source code of the MNDA pipeline in *R* is freely available on GitHub (https://github.com/H2020TranSYS/microbiome_dynamic). The LucKi cohort dataset is available upon request from the Euregional Microbiome Center (www.microbiomecenter.eu).

## Authors’ contributions

B.Y. and F.M. performed the main analysis and contributed to writing the text. G.G. performed the DMM analysis. N.V.B., M.M. and J.P. contributed in the LucKi cohort and helped interpreting the data and the results. J.P., B.S. and K.V.S supervised the project and contributed to writing the text.

## Acknowledgements

This study was embedded within the Euregional Microbiome Center (www.microbiomecenter.eu), a cross-border initiative on host-microbiome interactions between the University of Liège, Maastricht University, Maastricht University Medical Center+ and Uniklinik RWTH Aachen.

