## Additional File 1 for "Capturing the dynamics of microbiomes using individual-specific networks"

**Individual-specific network inference to capture dynamic microbiome interactions**

**Additional file 1:** **Global Network Construction**

MAGMA is the primary method for global network construction employed. It consists of several steps. Firstly, OTU’s abundances are modelled using a ZINB (zero-inflated negative binomial) that can also accept covariates. Furthermore, a maximum likelihood approach is used to infer the ZINB. On top of that a sparse precision matrix $\tilde{\theta}_{\rho}$ is estimated. Different values of penalization parameter $\rho$ yield different precision matrices, and hence $\rho$ is optimized considering three approaches as referred to in Cougoul et al. (2019). In this work we selected the rotation information criteria (RIC – Zhao et al. 2012) as default in the MAGMA package. Finally, the optimized penalization parameter $\rho^{*}$ is employed to identify the ASV network from the nonzero elements $\tilde{\theta}_{\rho^{*}}$. We refer the interested reader to the work of Cougoul et al. (2019) for an in-depth understanding of the MAGMA algorithm.

An alternative to MAGMA is the widely used SparCC approach. SparCC applies a log-transformation of the components and then estimates the Pearson correlations between those components. The algorithm calculates iteratively a “basis correlation”, with the underlying assumption that most pairs do not correlate. This approach reduces the amount of spurious negative correlations derived from the compositional nature of the data (Friedman et al. 2012). To amend the problem that zero would cause to the log transformation it employs a pseudo-count, i.e, assigning a small fraction to OTUs not detected in a sample. A fast implementation of SparCC, used in this study, is given by FastSpar.

There are many motivations behind MAGMA’s choice over SparCC. Cardinally, it can include covariates and does not need to adjust via pseudocounts. Moreover, MAGMA is able to infer direct associations using notions conditional independence. Notably, both SparCC and MAGMA work directly on the OTU counts. Hence no pre-transformation is needed. Moreover, both methods take into account the compositional zero-inflated and overdispersed nature of the data.
