## Additional File 2 for "Capturing the dynamics of microbiomes using individual-specific networks"

**Individual-specific network inference to capture dynamic microbiome interactions**

**Additional file 2: SparCC Analyses**

We have repeated the microbiome dynamic analyses with SparCC as global network construction method instead of MAGMA. The results are shown in Figure S10. We observe differences with MAGMA for the number and strength of connections, which appear to be more abundant and stronger in SparCC analysis. This can be explained by the lack of natural specification of the SparCC. Moreover, SparCC naturally yield a weighted network. It shows the impact the selected network construction method has on final results. Microbial co-occurrence networks generated via different inference methods have been shown to exhibit quite varying network properties (Kishore et al. 2020). The future will tell which methods are most accurate and robust and/or whether to consider ensemble approaches across mutiple co-occurrence networks in our framework.
