## Additional File 3 for "Capturing the dynamics of microbiomes using individual-specific networks"

**Individual-specific network inference to capture dynamic microbiome interactions**

**Additional file 3: Supplementary Figures and Table**

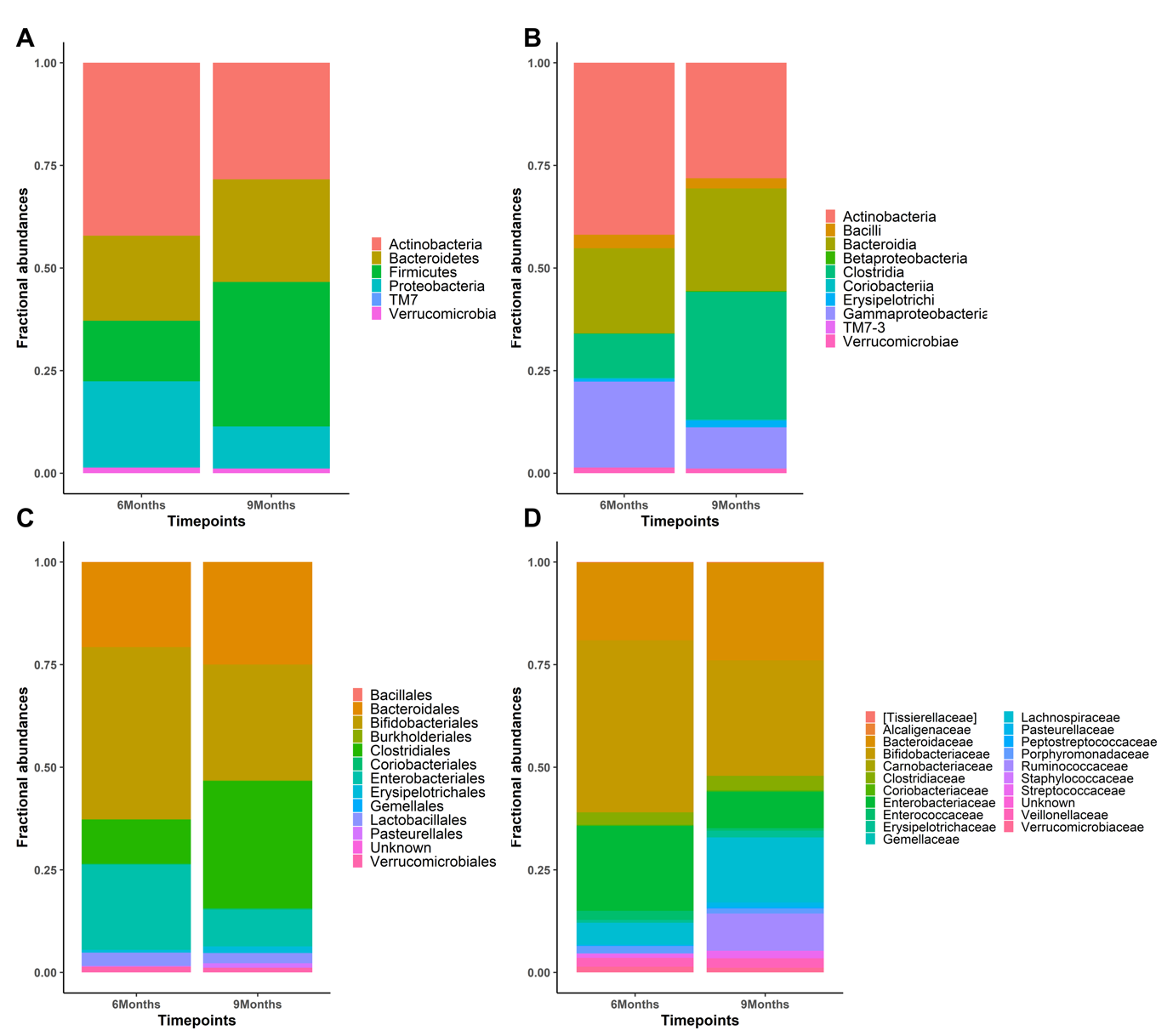

**Figure S1**: Fraction of relative abundances belonging to each (A) phylum, (B) order, (C) class, (D) family in 6 months and 9 months, on the 95 microbes selected with MAGMA pre-processing for the 69 paired newbors.

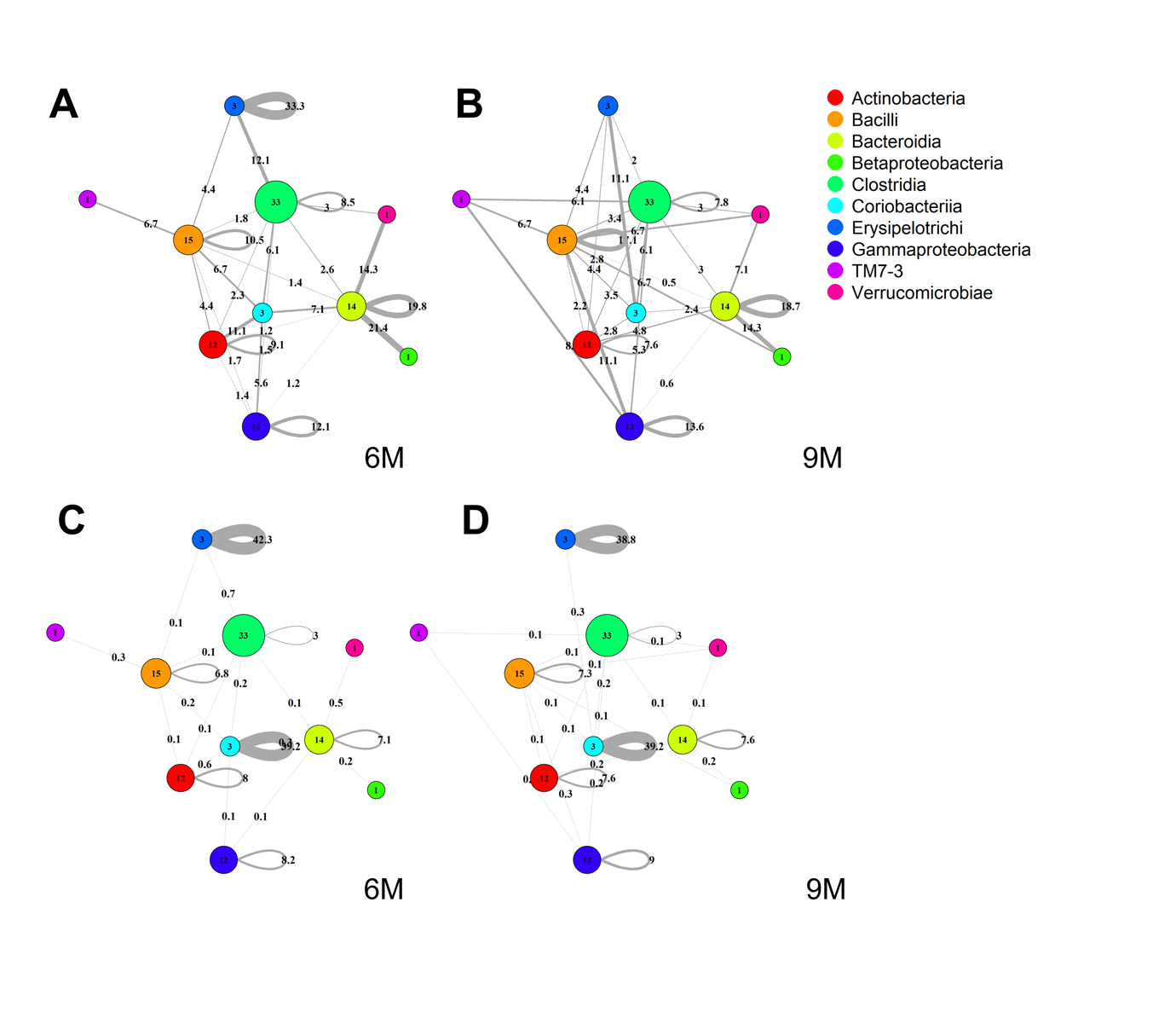

**Figure S2**: Representation of binary (top) and continuous (bottom) *Class* taxa interaction at month 6 (left) and 9 (right). From the binary graphs we can see an increase in significant interactions between taxa grouped per *Class* from month 6 to 9. This is not translated in weighted MAGMA networks.

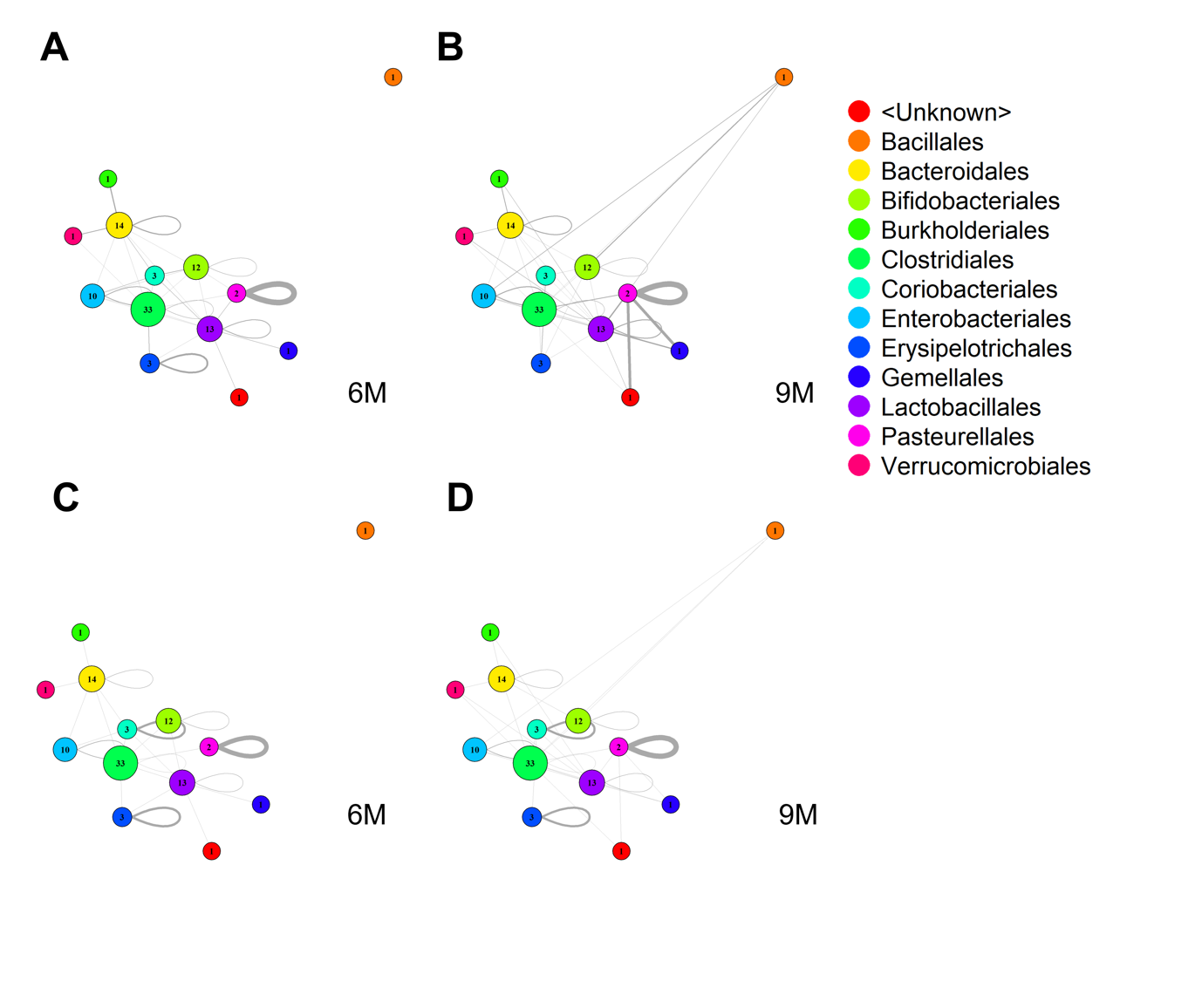

**Figure S3**: Representation of binary (top) and continuous (bottom) *Order* taxa interaction at month 6 (left) and 9 (right). From the binary graphs we can see an increasing of the significant interactions between taxa grouped per *Order* from month 6 to 9

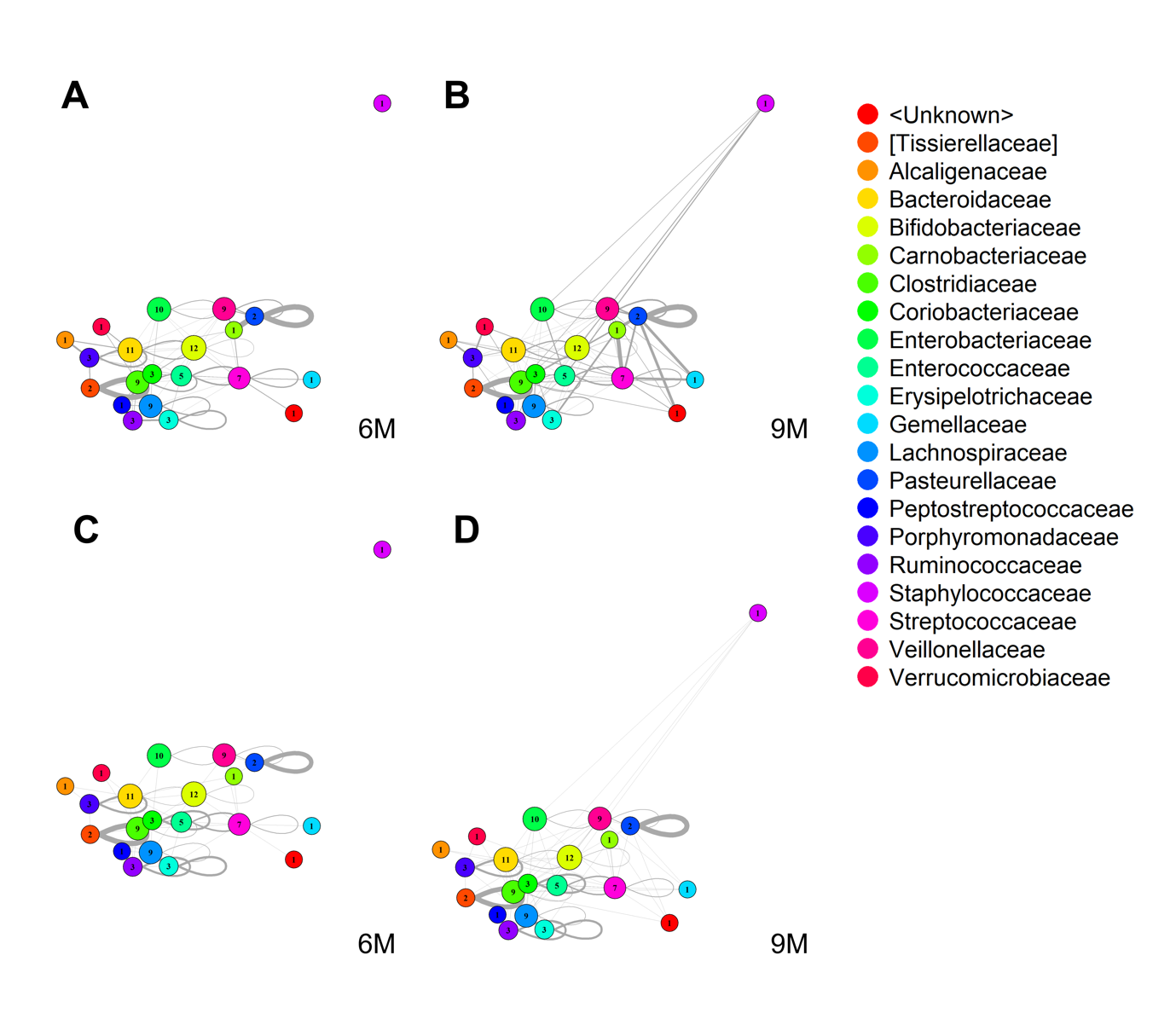

**Figure S4**: Representation of binary (top) and weighted (bottom) *family* taxa interactions at month 6 (left) and 9 (right). From the binary graphs we can see an increase in significant interactions between taxa grouped per *family* from month 6 to 9. Edge weights are not shown.

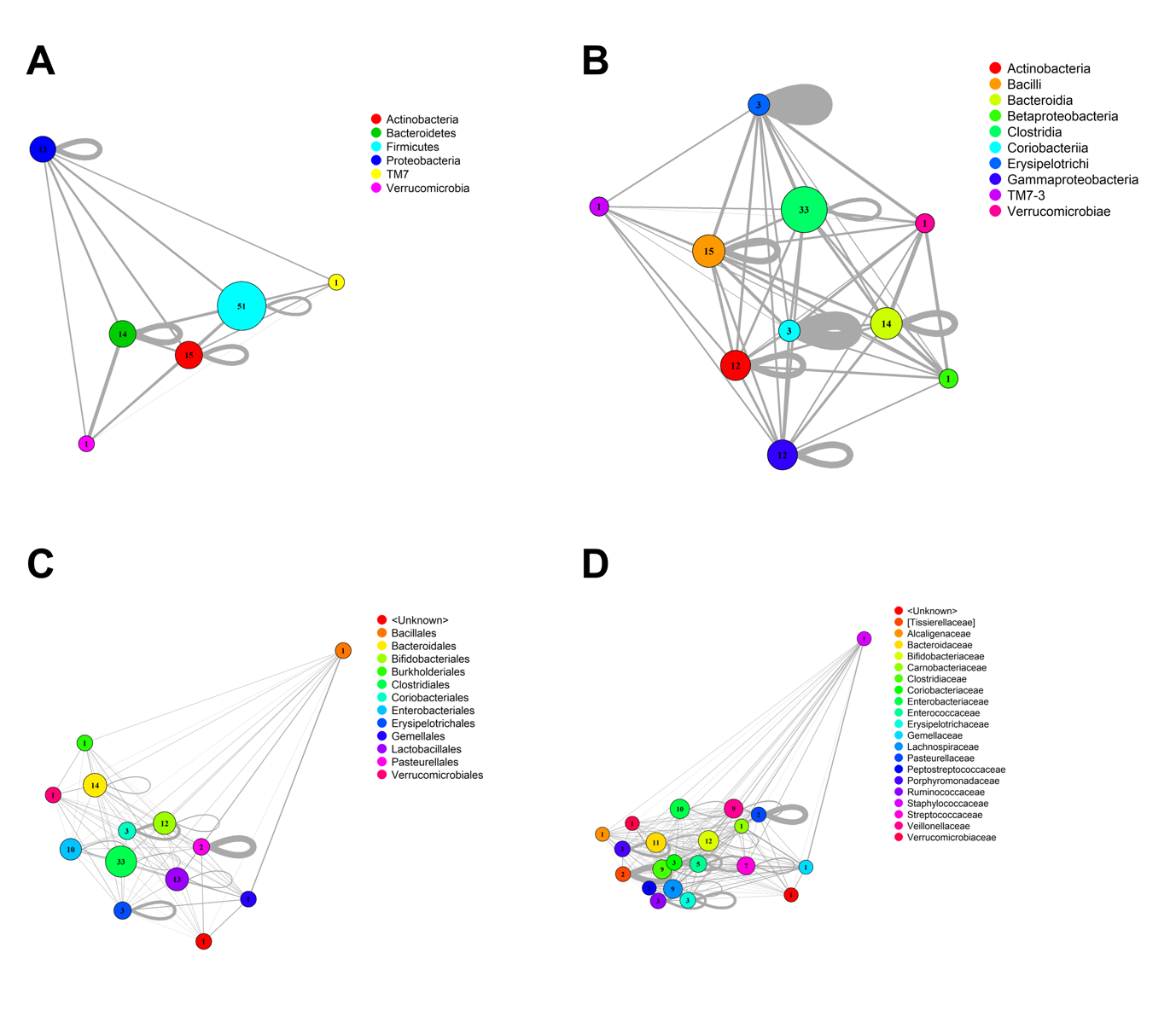

**Figure S5**: Representation of weighted taxa interactions at month 6, grouped for (A) Phylum, (B) Class, (C) and (D) Families with the SparCC method. SparCC does not sparsify as MAGMA

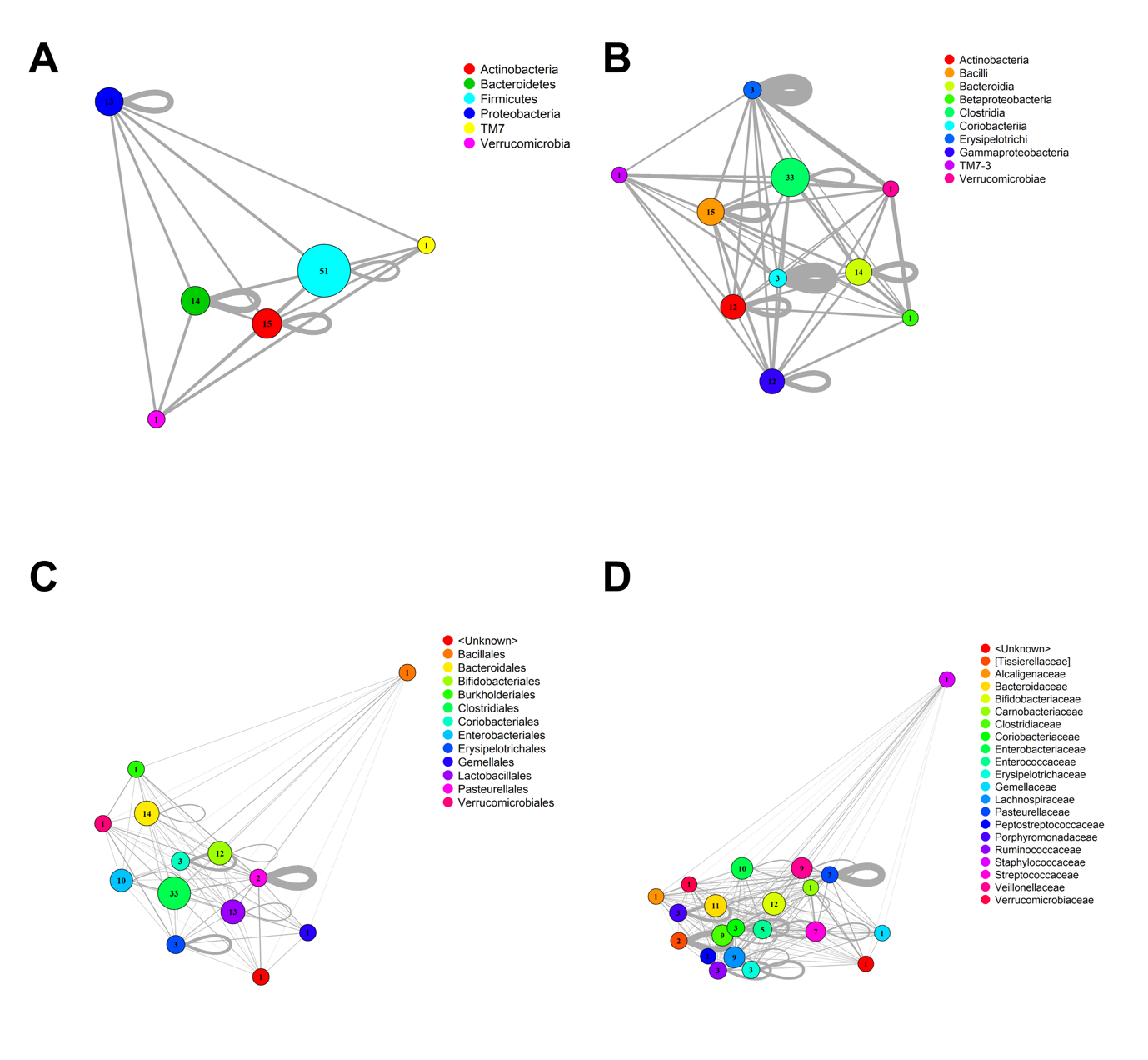

**Figure S6**: Representation of weighted taxa interactions at month 9, grouped for (A) Phylum, (B) Class, (C) and (D) Families with the SparCC method. SparCC does not sparsify as MAGMA

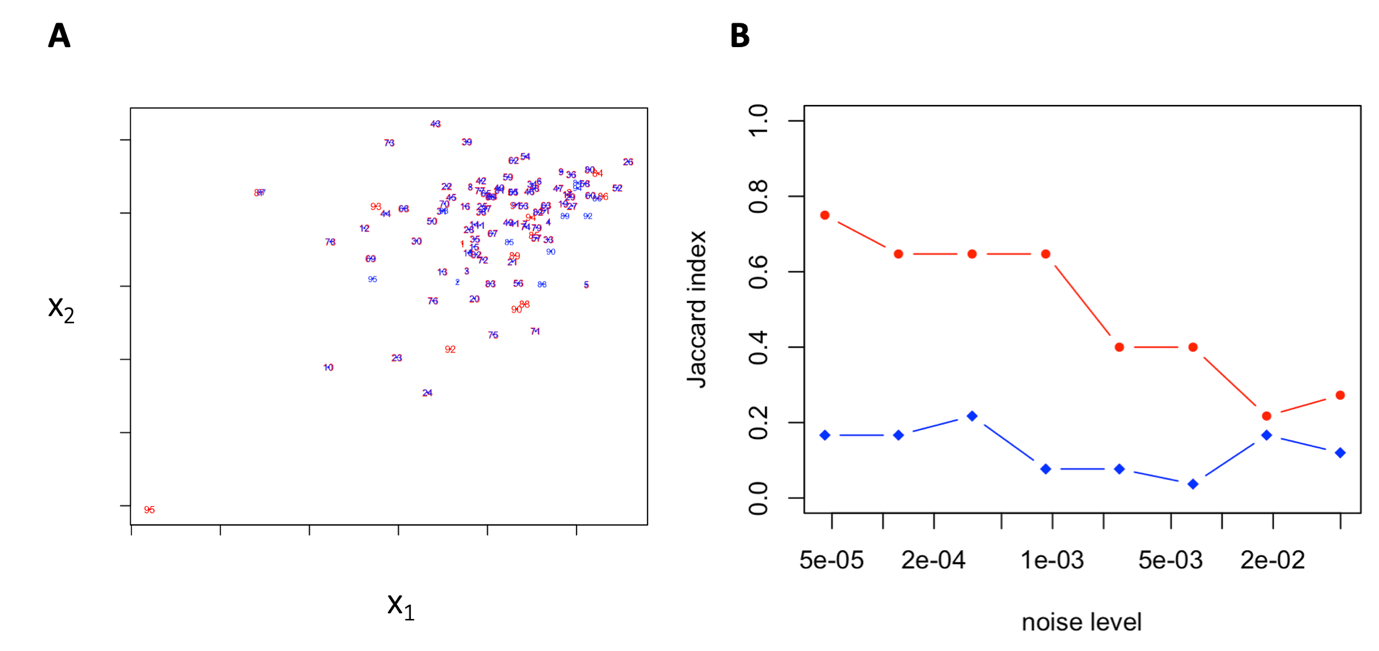

**Figure S7**: (A) A 2D representation of the EDNN embedding space for the simulated data. Each number is a node (red: layer 1 and blue: layer 2). As it can be seen, all the nodes whose neighbours are not changing fall on the top of each other except for a few that are defined to be different. (B) The Jaccard index between the pre-set varying nodes and the inferred varying nodes against the increase of the noise. The red and blue diagram correspond to the MNDA-based and eigen decomposition-based methods, respectively.

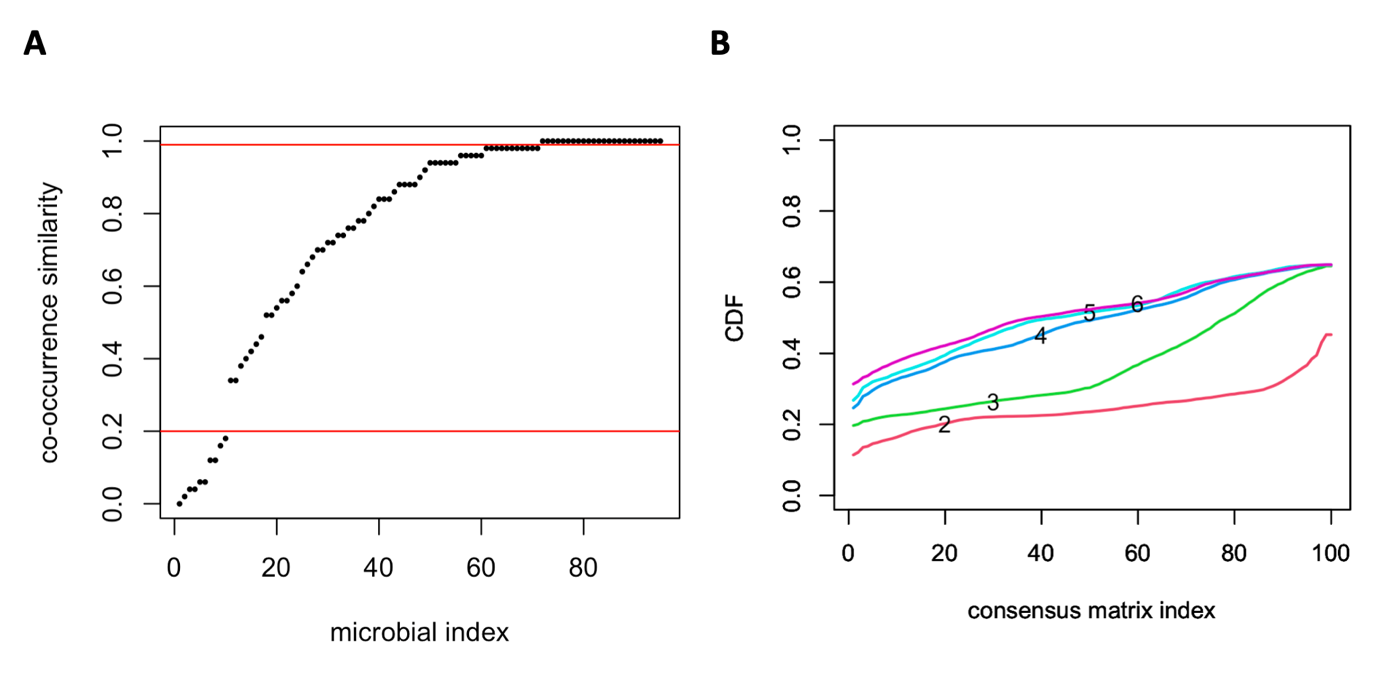
 **Figure S8:** (A) The selection of the highly and lowly dynamical microbes based on co-clustering similarities for each node pair, (B) The result of consensus clustering commutive distribution pointing to the existence of two robust clusters of microbes.

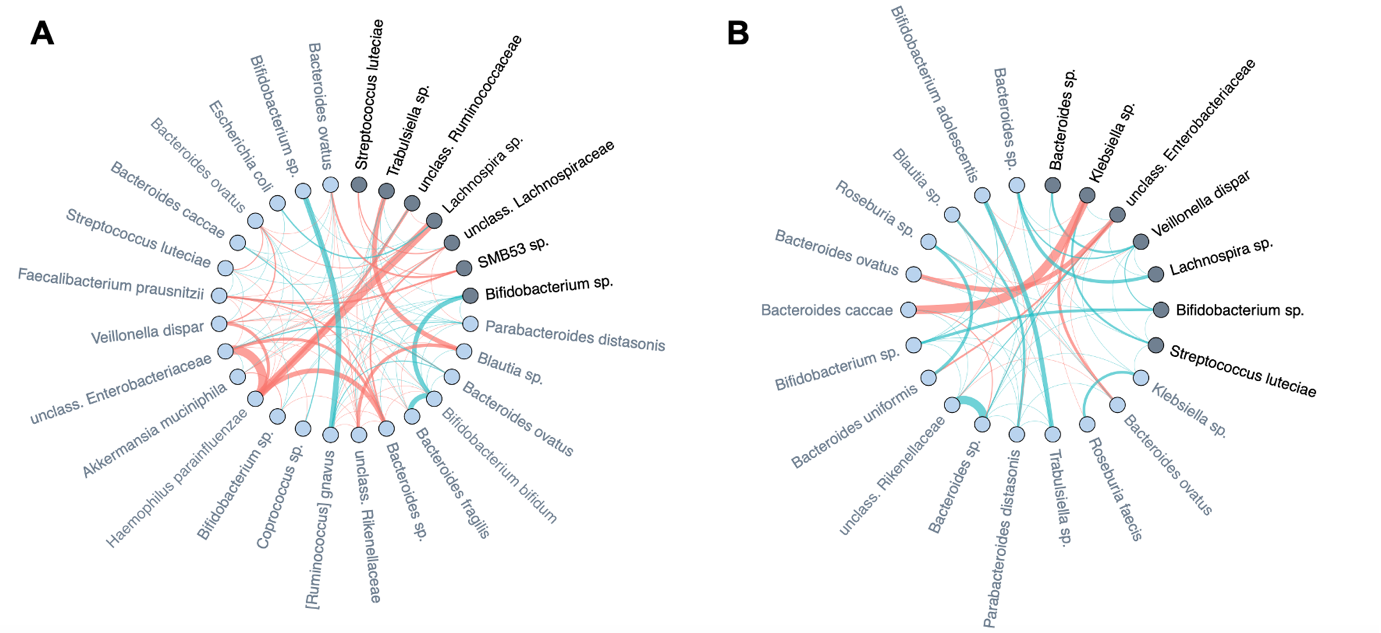

**Figure S9:** Comparison of the differences of averaged microbial co-expression networks restricted to the important microbes and their first level neighbours between month 6 and month 9 (Figure 10). (A) the difference network of c-section delivery subtracted from the difference network of vaginal delivery. (B) the difference network of persistent diet subtracted from the difference network of non-persistent diet.

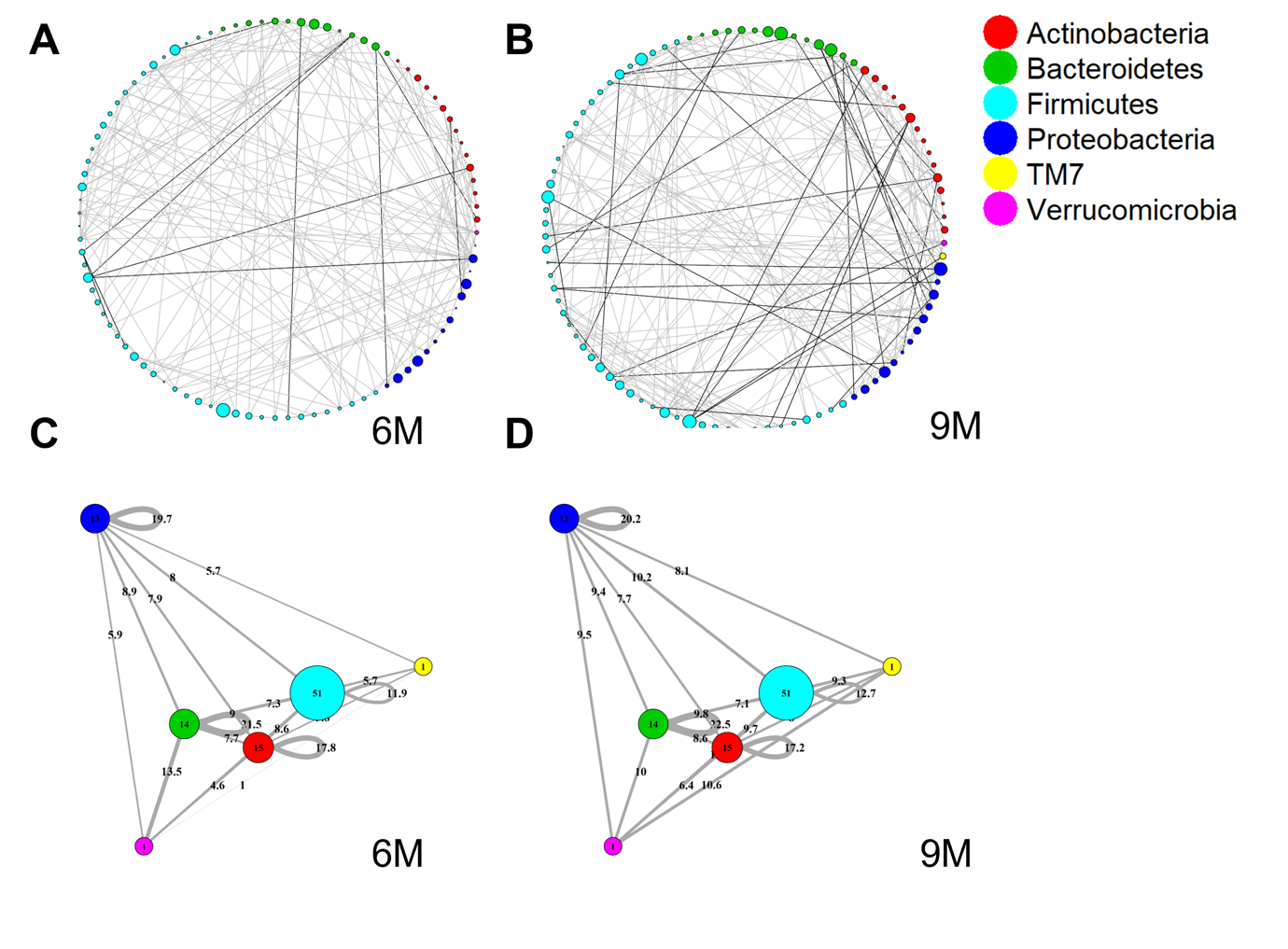

**Figure S10:** Global SparCC networks calculated on all subjects retained in the LucKi cohort at 6 (81) and 9 (74) months after birth. Color code corresponds to phylum classification. The thickness of an edge corresponds to the strength of association. Microbial taxa are organized on a circle according to phylum membership at time-point 6m (A) and 9m (B). In (C) and (D), respectively time-point 6m and 9m, edges are aggregated per phylum. Within- and across- phyla co-occurrences are computed via SparCC continuous networks at each time point. The size of nodes and thickness of edges correspond to phylum size and association strength.

**Table SI.** The list of microbes that swich their clusters from month 6 to month 9 with their corresponding genus-species names.

|  | **Microbe ASVs** |
| --- | --- |
| **Cluster 1 to cluster 2** | *Akkermansia muciniphila, Bacteroides fragilis, Bacteroides ovatus, Bifidobacterium sp., Bifidobacterium sp., Clostridium perfringens, Collinsella aerofaciens, Faecalibacterium prausnitzii, SMB53 sp., Streptococcus luteciae, Streptococcus sp., Streptococcus sp.* |
| **Cluster 2 to cluster 1** | *Bacteroides ovatus, Bacteroides sp., Bacteroides sp., Bifidobacterium adolescentis, Bifidobacterium longum, Enterobacteriaceae unclassified, Enterococcus sp., Escherichia coli, Faecalibacterium prausnitzii, Parabacteroides sp., Ruminococcus bromii, Ruminococcus gnavus* |
